# Oligomeric assemblies of plant biotin carboxylase revealed by cryo-EM and cross-linking

**DOI:** 10.64898/2025.12.30.697075

**Authors:** Hunter J. Madison, Luke Dunn, Youngki You, Gabriel Lemes Jorge, Ljiljana Paša-Tolić, Jay J. Thelen, Steven R. Van Doren, Adam L. Yokom

**Author notes:** Address correspondence to: Steven R. Van Doren, Adam L. Yokom.

## Abstract

Due to the interest in fatty acid synthesis by oilseed crops, we conducted structural studies of the biotin carboxylase (BC) subunit of the plastid acetyl-CoA carboxylase. Acetyl-CoA carboxylase catalyzes the first committed step in the fatty acid synthesis pathway and is highly regulated. Cryo-electron microscopy revealed that pennycress apo-BC forms a symmetric dimer and contains a subpopulation of a dimer-of-dimers. The domain of BC that closes over the catalytic cleft (the B-domain) appears to be dynamic, judging from the b-factors, normal mode analysis of BC structures, and its high susceptibility to acetylation. An increase in the BC concentration decreased the reactivity of the B-domain, however, suggesting structural hindrance. The partial protection of the B-domain was consistent with cross-links that formed between dimers of BC using a cross-linker cleavable in the mass spectrometer. Cross-links guided HADDOCK docking calculations suggesting a dimer of dimers of pennycress BC that is asymmetric, staggered, and tilted between dimers, with conservation in the interface. In contrast, a minimal population of a symmetric dimer of dimers with a small, non-conserved interface was observed by cryo-EM. Taken together, our structural models are the first for *Brassicaceae* family BC homologs and are the first from plants. These models suggest dimer interactions that might contribute to larger oligomers of BC and influence associations with other subunits of the heteromeric acetyl-CoA carboxylase.

## 1. Introduction

The oil content of harvested seeds is critical for food, biofuel, and animal feed, maintaining research interest in the *Brassicaceae* family of oilseed crops (1–4). The triacylglycerols stored in seeds and leaves in large part derive from fatty acids synthesized *de novo* in chloroplasts under substantial regulation. The initial committed step of fatty acid synthesis by autotrophs is catalyzed by acetyl-CoA carboxylase (ACCase), which combines acetyl-CoA with bicarbonate to form the malonyl-CoA required for cycles of fatty acid elongation (5, 6). The cytosol of eukaryotic cells contains a homomeric ACCase encoded by a single polypeptide chain (7, 8). However, bacteria and the plastids of most algae, dicots, non-graminaceous monocots express a heteromeric ACCase (het ACCase) with biotin carboxylase (BC) and carboxyl transferase (CT) catalytic subunits, connected functionally by the biotin carboxyl carrier protein (BCCP) (5–9).BC binds ATP, Mg^2+^, and bicarbonate, to form carboxyphosphate initially (6, 10). BC then condenses this activated carboxyl group with the biotin cofactor of BCCP (10). BCCP shuttles its carboxybiotin to the active site of CT (6, 11).

Key functional partners of BC in dicots include both BCCP1 and BCCP2 homologs (12–14) and three BCCP-mimicking, biotin attachment domain-containing subunits known as BADC1, BADC2, and BADC3. BADCs lack the loop which is biotinylated in BCCPs (13, 14). The occupancy of BC by BCCP and BADC subunits was proposed to change with the stromal pH of the day-night cycle (15). BADC1 and BADC3 appear to be sequestered by PII for joint regulation of C and N metabolism in Arabidopsis (16). Consequently, BC in plants and algae can be viewed as a hub of protein-protein interactions which regulate BC and in turn ACCase activity (13). We sought to characterize the BC from the *Brassica* species *Thlaspi arvense*, i.e. field pennycress, because pennycress is a prospective winter cover crop for oilseeds. Efforts to engineer pennycress take advantage of its diploid genome for modifications to enhance oil and the distribution of fatty acids (4).

Experimental structures of BC subunits from several bacterial species are available. High-resolution crystal structures of BC from *E. coli* ACCase are available in the apo form (17) and multiple complexes with substrates and ADP, ATP, or non-hydrolyzable nucleotide (18, 19, 10, 20, 21). Experimental BC structures are available from three thermophilic bacterial species (19, 22, 23) and pathogenic bacterial species as well (24–26). These studies revealed that BC comprises three domains, an N-terminal or A domain, a flexible B domain, and a C-terminal or C domain (Fig 1A) (17). Together, the A and C domains are arranged in an ATP grasp fold (6). Most of the structures of BC from bacterial ACCases are dimeric, though a tetramer was observed with BCCP bound (23). The crystal structures of BC from the green microalga *Ankistrodesmus* sp form similar dimers (27). Up to six BC-containing chains were observed in oligomers of propionyl-CoA carboxylase (28). Oligomers of BC, which included BCCP, of at least 800 kDa were found in the soybean chloroplast stroma (9). BCCP-containing BC oligomers, prepared from recombinant Arabidopsis subunits of up to 1 MDa were observed (29). Tubular filaments of hetACCase from *E. coli* were observed by cryo-EM (30). Despite the many structures of bacterial BC, there have been no experimental structures of a plastid ACCase from plants to our knowledge. Which oligomeric states to expect has been unclear. Experimental structures are also needed for understanding associations of subunits in het ACCase.

**Figure 1.**
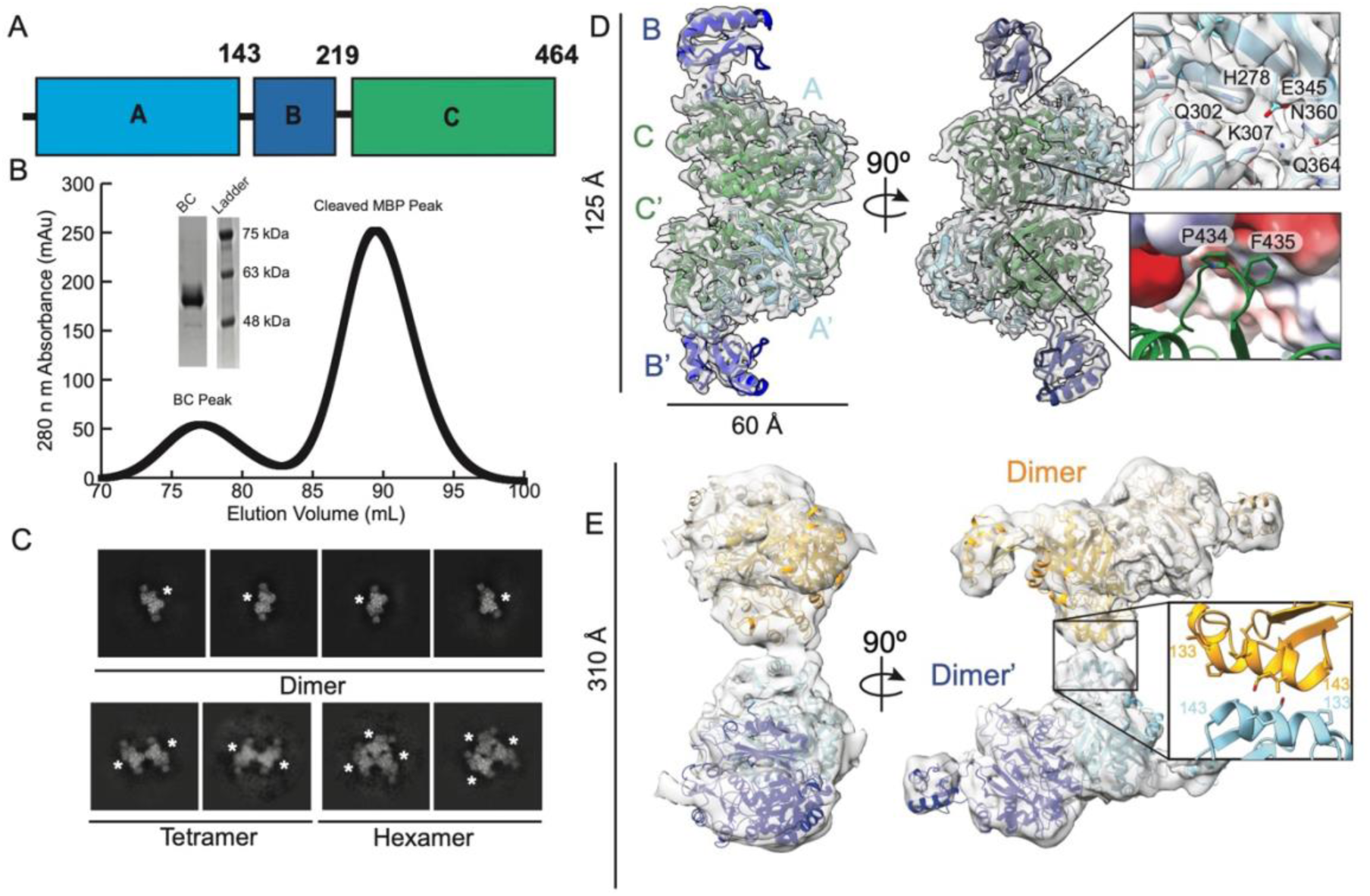
Cryo-electron microscopic analysis of pennycress BC. A) Domain schematic of pennycress biotin carboxylase. B) Size exclusion chromatography shows a purified peak for pennycress BC (peak 1) along with TEV cleaved MBP (peak 2). Inset shows corresponding SDS-PAGE for the BC peak. C) Representative 2D class averages for dimer, dimer of dimers and hexameric BC are shown. White asterisks mark each BC dimer. D) Final sharpened cryo electron microscopy map for the BC dimer with a nominal resolution of 2.85Å. Model is colored BC domain structure found in panel A. Top inset - close up of active site EM density with residues labeled. Bottom inlet - close up of the dimer interface with β-turn (aa 429-440) shown in green and opposing dimer interface shown as an electrostatic surface. E) 8 Å cryoEM map for dimer of dimers BC particles shown docked with two pennycress BC dimers. Models are denoted BC and BC’ and colored as shades of orange and blue, respectively. Inset – interaction of α-helix containing residues 132-143 with residue labels.

We report the first cryo-EM structures of a plant BC (from pennycress) without ligands, revealing a predominant dimeric form, and a low population of a symmetric dimer of dimers. Structural bioinformatics and surface reactivity suggested the lid domain to be dynamic, though a little less accessible at higher concentrations of BC. Finally, cross-linking mass spectrometry on BC samples suggest an asymmetric dimer of dimers with a larger interaction interface than the symmetric dimer of dimers observed via cryo-EM. Taken together, our data suggest that plastid BC samples multiple oligomeric states, which could be central to assembly and function of het ACCase complexes in plants.

## 2. Results

### 2.1. Structure of pennycress BC

To investigate the structural organization of BC from a strategic plant, we aimed to purify recombinant pennycress BC. Affinity purification via an engineered N-terminal Maltose Binding Protein (MBP) tag yielded soluble BC. Subsequent protease cleavage was used to remove the MBP tag. Notably, BC remained soluble without the presence of the MBP tag. Further purification showed a monodispersed Size Exclusion Chromatography peak and a single band via SDS-PAGE gel analysis (Fig 1B). Purified BC was concentrated to 13 µM (0.7 mg/mL) and vitrified for structure determination via cryo electron microscopy. Initial screening data showed clear 2D class averages with a two-fold axis of symmetry, similar to the previously published BC structures from *E. coli*. Iterative 2D classification was used to remove poor quality particles. During final rounds of 2D classification, three distinct oligomeric states were observed. The majority of the 2D averages contained a dimeric BC density while a small population (< 1% of total particles) contained a dimer of dimers or hexametric BC oligomers (Fig 1C). This was unexpected given the monodispersed peak observed in size exclusion chromatography (Fig 1B). We suspect the tetramer and hexamer architectures may represent transient states.

From these data we obtained the first 3D model for pennycress BC dimer at a nominal resolution of 2.85 Å (Fig 1D, S1 and Table S1). Two monomers of pennycress BC readily fit into our map and are denoted as BC and BC′ (Fig 1D). The pennycress BC dimer is analogous to published bacterial BC structures (20, 21, 31–36); see section 2.2. The A and C domains, defined in ref (17), and the interface between the dimers have the highest overall resolutions (<3 Å) (Fig 1D, insets). The peripheral regions including the B domain exhibited slightly lower resolution (>3.5 Å), consistent with dynamic flexibility expected in this region (Fig S1).

Evolutionary Trace analysis of BC sequences identified a number of core residues which are conserved across organisms (Table S3). Among these are a broad active site formed by the C and A domains (Fig S2A). In our pennycress BC dimer, clear density is present for these conserved residues. This includes the BC bicarbonate binding loop (R362, Q364, V365, E366, R408) and ATP binding sites (Q302, E345, L347, N360, T508) (Fig 1D, top inset). Furthermore, there is an extensive dimer interface between the BC and BC’ monomers via 1406 Å^2^ of buried surface area. A β-turn (aa 429-440) from each BC monomer extends into a binding pocket of the opposing BC (Fig 1D, bottom inset). The core of the dimer interface is highly conserved within our Evolutionary Trace data (Fig S2B). These features are found in prior *E. coli* BC structures (17) and confirm that the pennycress BC structure mimics those models.

We were unable to obtain a 3D reconstruction for the hexameric state due to the limited particle count and preferred orientation of these particles. The dimer of dimers resolved to a moderate resolution map of ∼8Å (Fig S1). Docking of two copies of our high-resolution BC dimer model revealed a novel interface between adjacent A domains (Fig 1e, inlet). A single α-helix containing residues 132-143 forms the small interface (193 Å^2^) between dimers in a ‘kissing’ arrangement (Fig 1E). Our pennycress BC dimer-of-dimers architecture is structurally distinct from the published *E. coli* BC:BCCP tetramer (PDB:4HR7). In that structure, binding of an *E. coli* BCCP occurs at the homologous α-helix (aa 64-73) (37). However, the tetramerization of the BC from *Chloroflexus auranticus* via a N-terminal swapped BCCP domain (PDB:8HZ4) (38) is analogous to the “kissing” dimer-of-dimers observed for pennycress BC. Overall, our structural characterization of pennycress BC revealed the first models for two states. These data suggest that there are transient but distinct oligomeric states present.

### 2.2 BC structures include dynamic and closable B domain lid

The B-factors are elevated within the B-domain of pennycress BC, as well as in loops of the A and C domains flanking the active site (Fig 2A). The working hypothesis that the lid and flanks of the active site are dynamic was followed up by structural bioinformatics comparison with ten bacterial BC structures. The sequence identity of these enzymes with pennycress BC ranged from 51.8 to 54.5% (Fig S3A). A core of 420 amino acids shared by the BC structures was identified using the Bio3D package (39). Residues in this core that moved in a highly correlated fashion had a high atomic movement similarity matrix (40). This identified the A domain moving jointly, the small B domain, and the large C domain (Fig 2B). Further inspection of the correlation matrix suggested a relatively rigid C_1_ subdomain providing a floor to the active site (blue in Fig 2C) followed by a more mobile C_2_ subdomain (marine blue in Fig 2C).

**Figure 2.**
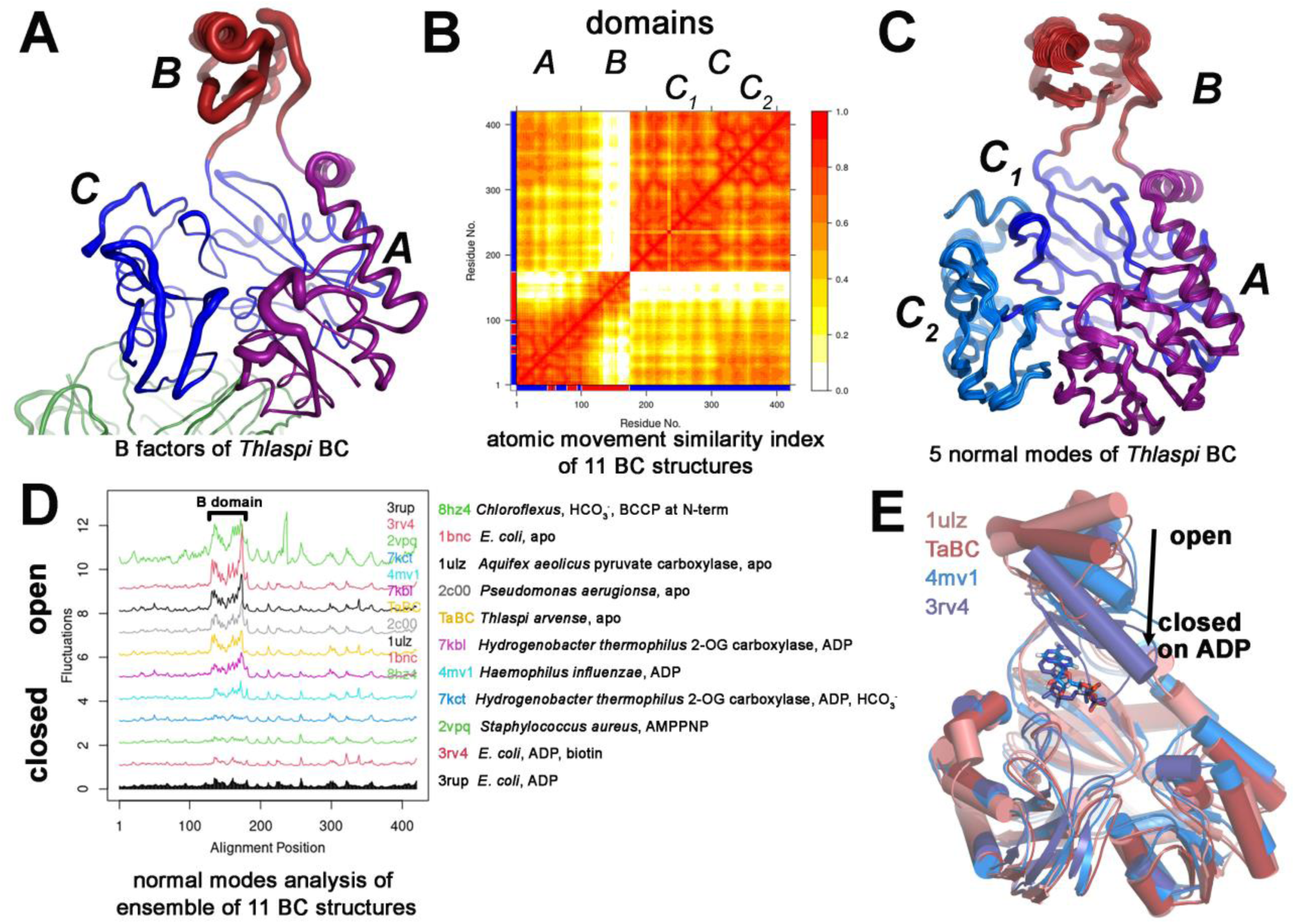
Movable domains of BC catalytic subunits from 11 species compared using structural bioinformatics. A) The diameter of the tube is proportional to the B-factors in the cryo-EM structure of BC from *T. arvense*. The A, B, and C domains are colored purple, red, and blue, respectively. B) BC structural from *T. arvense* and ten closest orthologues, all bacterial, were fitted to a shared structural core of 420 residues by Bio3D. The GeoStaS algorithm (40) computed the atomic movement similarity index among the 420 residues shared by the structures. This corroborated the recognized domains and movement of the B domain (31), and suggested perceptible subdomains within the C domain. C) Normal modes analysis (NMA) was performed on the ensemble of 11 structures using Bio3D (62, 63). A bundle of the 55 structural frames demarcating the simulated structures of *T. arvense* BC is plotted. The color scheme is shared with panel A but for the C1 and C2 subdomains in two shades of blue. D) The sizes of the fluctuations simulated by NMA are plotted vs. shared residue number of the 11 structures. The apo models with open active sites are plotted above while the structures closed upon the nucleotide in the active site are plotted below. The legend to the PDB accession codes is given to the right. E) PCA of the 11 structures grouped the cryo-EM model of pennycress BC with the three crystal structures in the dendrogram of Fig. S4B and plotted here.

Normal modes analysis (NMA) (39) of the group of 11 structures predicted and compared their dynamics (Fig 2D). This corroborated the mobility of the B domains of the five open, apo structures, including our pennycress BC dimer (Fig 2C,D). The mobile B domain (residues 127-178) was visible in the core 420 residues and is marked on Fig 2D. Potential directions of rotation of the B domain are suggested by five normal modes of *T. arvense* BC simulated in Video S1. The NMA suggests additional mobility within the A and C domains (Video S1). The mobility of the B domain was minimized in four structures with this lid domain closed in the nucleotide bound state, i.e. *E. coli* BC with ADP, the BC of *H. thermophilus* 2-oxoglutarate carboxylase, and *S. aureus* BC bound to ATP analog (Fig 2D). Two ADP-bound BC structures from *H. influenzae* and *H. thermophilus* appeared to have intermediate mobility of the B domain. Principal component analysis followed by clustering suggests the pennycress BC dimer to be most similar to the BC structures of apo *A. aeolicus* pyruvate carboxylase, *H. influenzae* with ADP bound, and *E. coli* with both ADP and bicarbonate bound (PDB: 1ULZ, 4MV1, and 3RV4, respectively, in Fig S3B). Structural overlay of these four structures highlights the rotation that closes the B domain to bind the nucleotide (Fig 2E). The top three modes were rotations of the B domain in these structures. This rotation of the B domain acting as a lid seems intrinsic to capture ATP and release of ADP. The top normal mode of each BC structure analyzed resembled the closing of the B domain lid upon nucleotide binding (Video S1, Fig 2E). The *E. coli* structure had a fourth rotation of the B domain. BC structures had normal modes rotating the A and C domains among the top five modes (see frames 34 – 55 of Video S1). Our NMA highlights that pennycress BC behaves similarly to other BC homologs, specifically the B domain rotations which have been seen in other structural models.

### 2.3 High susceptibility of B domain lid to acetylation

We sought an experimental appraisal of BC dynamics and interactions beyond comparison to structural homologs. Surface accessibility was tested by an acetylation assay (41, 42). Briefly, BC was incubated with *N*-succinimidyl acetate (NHS-acetate) and digested using endoproteinase GluC to generate labeled proteolytic fragments with acetylation of lysine residues. Peptides detected reproducibly covered 21 of the 22 Lysine residues (the exception being K74) in the mature sequence of BC (Fig. 3A). Further replicates were performed to test the dependencies of acetylation on NHS-acetate (41, 42) and BC concentrations. Acetylation was detectable at 20 Lysine residues, as well as at Y268 and S454. Among the seven most reactive residues (Fig. 3A), four reside in the exposed B domain (K228, K249, Y268, and K271) along with K186 nearby in the A domain (Fig. 3D,E). Nine residues showed intermediate reactivity; K107 and K113 in the A domain, K255 in B domain, and K287, K416, S454, K502, K505, and K513 in the C domain (Fig. 3A,D,E). Poor reactivity occurred at the remaining seven lysine residues (K194 in the A domain and K307, K323, K458, K474, K493, and K528 in the C domain between K194 of the A domain to D205 of the B domain may limit the range of dynamics of the B domain. Nonetheless, the highest acetylation levels are found in the B domain supporting the NMA hypothesis that the B domain of pennycress BC is dynamic in the apo state (Fig. 2A-D).

**Figure 3.**
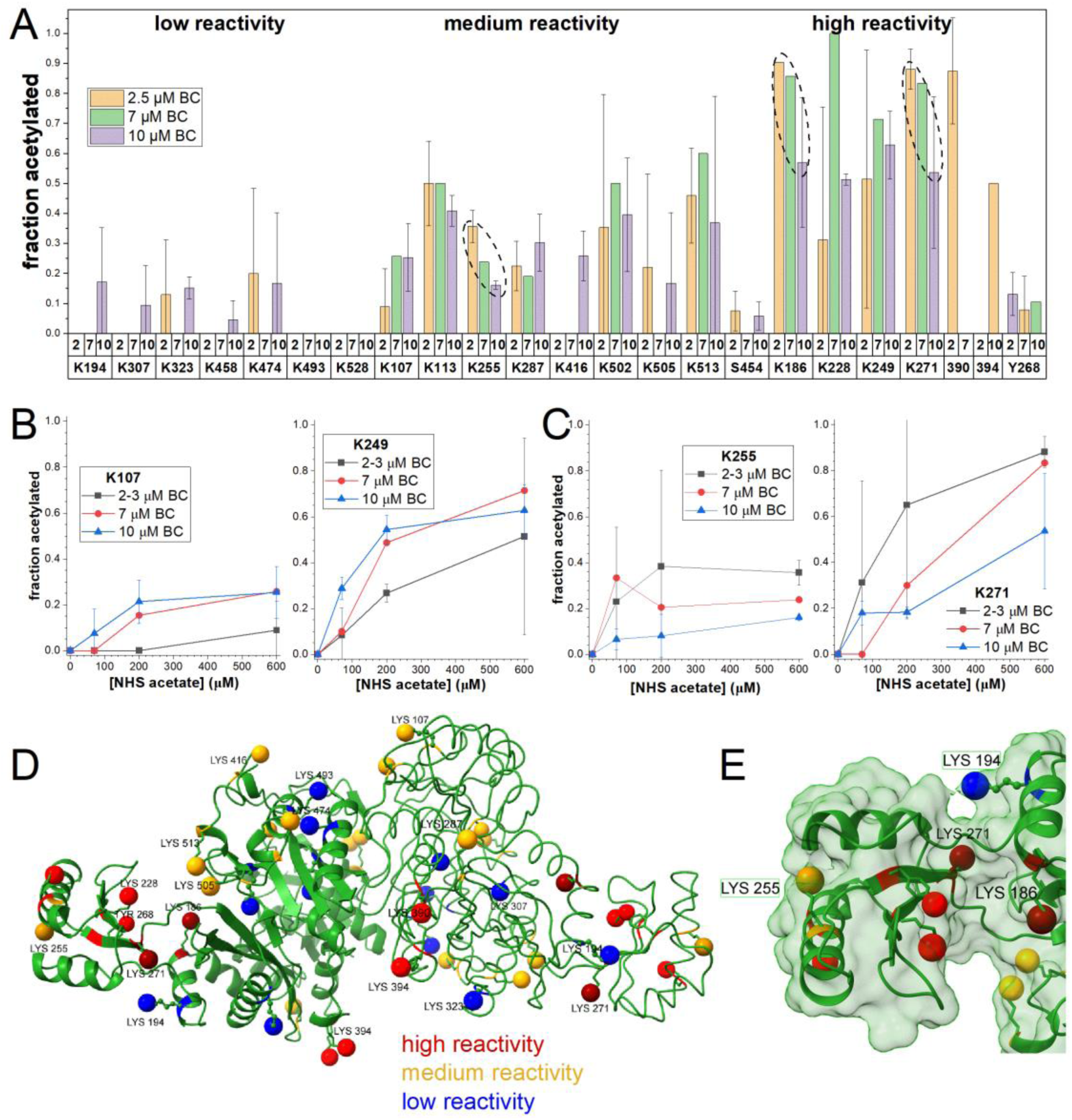
Surface accessibility of the catalytic subunit BC mapped by protein painting, i.e. acetylation of lysine residues. The fraction of each side chain acetylated by NHS-acetate was calculated as the proportion of Glu-C-generated peptide fragments modified. The series of reactions were repeated twice at 2 to 3 μM BC, once at 7 μM BC, and twice at 10 μM BC (Fig. 3B,C). Acetylation of all lysine residues and two hydroxyl-containing sidechains was quantified. *A*, fraction acetylation of these sidechains at 600 μM NHS-acetate (*N-*succinimidyl acetate) and [BC] of 2 – 3 μM (n=2), 7 μM (n=1), and 10 μM (n=2). The ellipses with dashed lines point out cases of acetylation appearing to decrease with increased [BC]. *B*, examples of concentration dependencies in which acetylation tend to be enhanced at higher [BC]. *C*, examples of concentration dependencies in which acetylation tended to lessen at higher [BC]. *D*, sites of acetylation are marked on cryo-EM-derived coordinates the dimer of BC were colored by reactivity. The darker shade of red (K186 and K271) or gold (K255) denote decreased reactivity at higher [BC]; see panels A and C. *E*, close-up view of residues near the highly exposed B domain that were protected from acetylation in general (K194) or at higher [BC] (K186, K255, K271).

However, K255 on the apex of the B domain lid and K186 and K271 in the interface between A and B domains become partly protected from acetylation in reactions with the higher [BC] concentration (10 μM). This raised the question of how higher [BC] can decrease dynamics around the active site. We pursued cross-linking mass spectroscopy to investigate BC oligomeric structure at concentrations beyond the capabilities of cryo-EM.

### 2.4. DBSU cross-links suggest an asymmetric dimer of dimers

SDS-PAGE analysis of DBSU crosslinked BC (at 10 μM BC) showed bands at dimeric and tetrameric molecular weights (Fig. S4). Mass spectrometry analysis revealed eight cross-links that were present within the monomeric structure of BC. Additionally, eight cross-links could not be explained by interactions in the monomer or dimer structures of BC (Tables 1 and S4). Additionally, these cross-links could not be rationalized via our cryo-EM symmetric dimer of dimers model (Fig 1E).

**Table 1.** Crosslinks that could not be mapped within the dimer of BC.

| Crosslink<sup>A</sup>, Lys-Lys |
| --- |
| 186 - 186 |
| 186 - 390 |
| 186 - 394 |
| 186 - 416 |
| 307 - 390 |
| 307 - 394 |
| 390 - 390 |
| 394 - 416 |

<sup>A</sup> The crosslinks were formed with the MS-cleavable disuccinimidyl dibutyric urea (DSBU), which spans 12.5 Å between its carbonyl termini.

Therefore, we utilized cross-link-guided docking using HADDOCK 2.4 (43) to determine tetrameric ensembles of BC to explain these cross links. A first phase of HADDOCK calculations on tetrameric BC identified four mutually compatible cross-links across the BC dimers (Fig. 4A, red-labeled Lysine residues). These were K390 and K394 of one dimer (green) to K186 and K307 of the second dimer (purple). A second phase of structural modeling using crosslinked-based distance restraints implicated three additional lysine cross-links on the periphery of the dimer-dimer interface. These extra lysine crosslinks are compatible with the original core of four cross-links identified in the first phase. These peripheral interchain cross-links are formed from K390 to K390, K416 to K186, and K416 to K394 (Fig. 4A, light blue labels).

**Figure 4.**
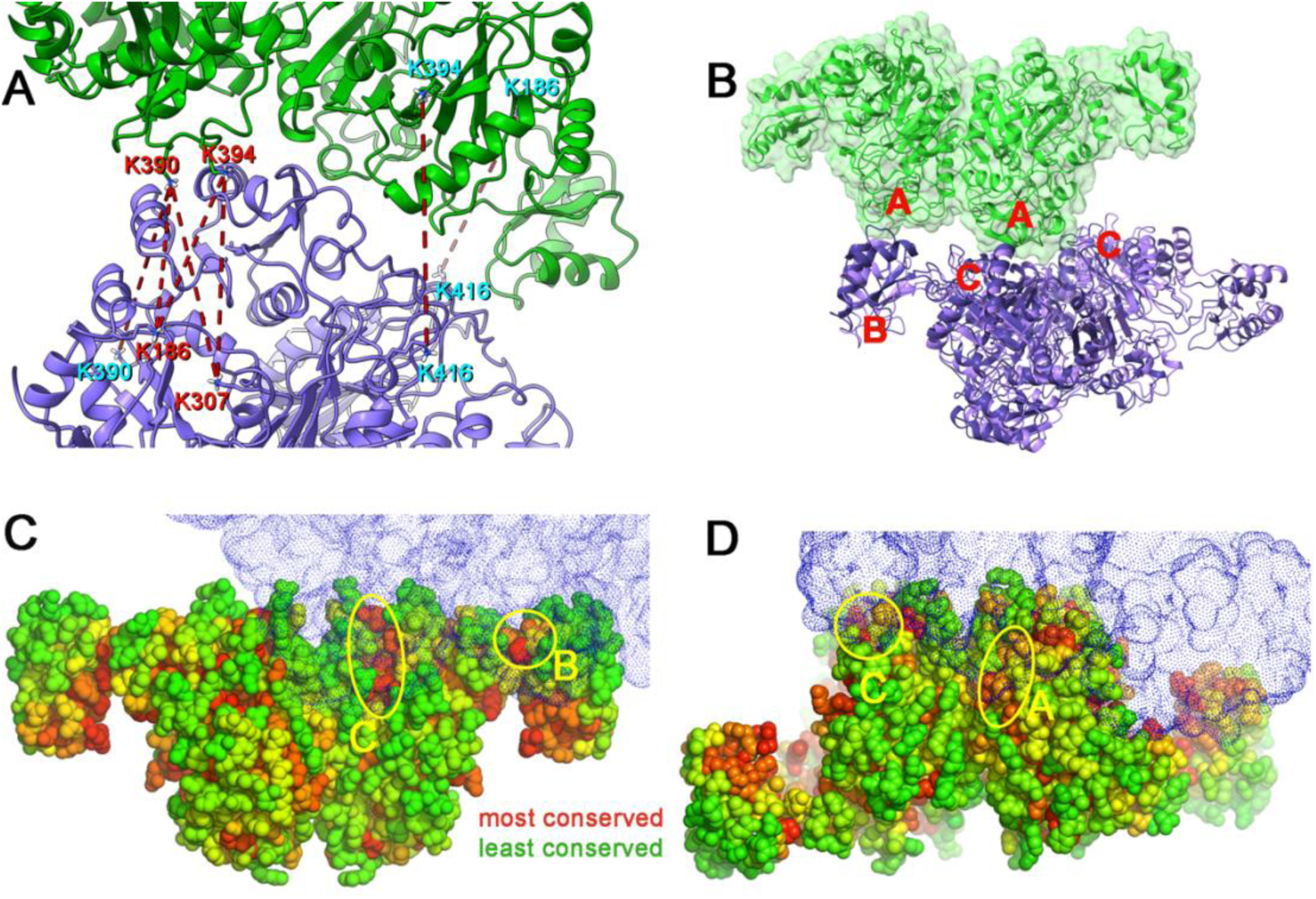
MS-detected crosslinks implicated an asymmetric dimer of dimers, with an A domain contacting either the C domains or B domain of the other dimer. (A) The four core cross-links are labeled with red text and marked on structural coordinates from HADDOCK calculations constrained by all seven cross-links. The dimers are colored green and purple respectively. (B) The domains in contact are labeled with letters. A structure from cluster 1 is plotted with a darker purple ribbon than the lighter purple of a structure from cluster 2. (C, D) The surface conservation calculated by Evolutionary Trace analysis colors the most conserved residues red, the next most conserved orange, and on to the least conserved residues in green. Conserved patches in the dimer-dimer interface are marked with yellow ellipses and the domain. The plotting of one dimer with purple dots on its accessible surface allows viewing of its binding surfaces on the other dimer.

The HADDOCK ensemble of BC dimer of dimer structures contained two similar clusters (Table 2, Video S2). Three models grouped closely into cluster 1 and four models in cluster 2. Cluster 1 buried ∼240 Å^2^ more surface area between the dimers when compared to cluster 2 (Table 2). There are a number of subtle changes in the BC dimer position including offset, tilt and proximity of the dimers in cluster 1 and 2. The angle of crossing between the two dimers is 11° larger in cluster 1 than cluster 2. The offset of the two dimers along the long axis of one dimer averaged about 4 Å longer in cluster 2 than cluster 1 (Table 2). These shifts suggest the degree of uncertainty in the positioning of the dimer-dimer interface consistent with seven crosslinks and contours of the interface.

**Table 2.** Positioning of the two dimers in the two clusters of structural solutions.

| Cluster | Surface area buried in dimer-dimer interface, Å<sup>2</sup> | Offset along long axis, Å | Tilt between long axes of dimers<sup>A</sup> | Closest approach between long axes<sup>A</sup>, Å |
| --- | --- | --- | --- | --- |
| 1 | 1226 ± 36 | 22.3 ± 0.4 | 33 ± 3° | 44.7 ± 0.8 |
| 2 | 986 ± 94 | 26.0 ± 1.6 | 22 ± 1° | 46.2 ± 0.7 |

<sup>A</sup> The tilt and closest approach were measured between longitudinal axes drawn from Gln253 to Gln253 in the B domains of each dimer.

One BC dimer is translated along the long axis by 22 Å or more, rotated by roughly 100°, and tilted by at least 22° relative to the other BC dimer (Table 2, Fig. 4B). The A domain of one BC dimer (green) asymmetrically fills the groove between the C domains of another BC dimer (purple; Fig. 4B). Additionally, the other A domain (green) contacts the back of B domain of the other dimer (purple). This contact may limit the opening of the bound B domain and thereby account for the decreases in reactivity in our acetylation assay at 10 μM BC (Fig. 3A,C). In view of reports of larger oligomers of BC (9, 29), it is conceivable that the associations of BC dimers observed could be repeated to form larger oligomers. To investigate this further, we returned to the Evolutionary Trace data (Table S3). Residues at the back of the B and C domains on BC dimer are highly conserved. Their presence within the dimer-dimer interface supports this dimer of dimers (Fig. 4C,D). Conservation on the A domain within the dimer-dimer interface is more moderate (Fig 4D). We believe the dimers of dimers from crosslinking mass spectrometry, HADDOCK modeling, and cryo-EM represent unique assemblies at varied concentrations of BC.

## 3. Future perspective

Cryo-EM and cross-linking approaches found the biotin carboxylase from pennycress to adopt multiple oligomeric states, including dimers, dimers of dimers, and hexamers.

This raises the question of oligomeric states of BC regulating activity, stability, or assembly of het ACCase complexes. Possible relationships of BC oligomeric states to interactions with molecular partners such as other ACCase subunits (CT, BCCP isoforms, and BADC isoforms) and substrates (ATP, bicarbonate, and biotin) could prove important. It is unclear whether the oligomeric states of BC observed are specific to plants or could generalize to BC of other taxa. Future translational studies can utilize the structural data presented here to test effects of BC oligomerization on seed oil production.

## 4. Materials and Methods

### 4.1. Purification of BC

#### Constructs

The BC construct for recombinant protein expression was generated by subcloning the mature pennycress BC sequence (entry CAH2072050.1 in BioProject PRJEB46635) into a construct containing an MBP-TEV epitope tag using standard molecular biology techniques. TargetP - 2.0 (44) predicted the chloroplast transit sequence to comprise the first 57 amino acids. The sequence is listed in Supplemental Tables S3 and S4.

#### Protein expression and purification

Expression was performed in *Escherichia coli* BL21(DE3) cells. BL21(DE3) cells were cultured in Luria Broth medium with appropriate antibiotics at 37°C until an optical density of 0.6 was achieved. The growth temperature was lowered to 16°C, and protein expression was induced by adding isopropyl β-D-thiogalactopyranoside (IPTG) to a final concentration of 1 mM. Expression was carried out for 18 hours before pelleting the cells via centrifugation at 4 °C for 30 min, 3500g. The pellet was resuspended in lysis buffer containing 50 mM Tris pH 7.5, 150 mM NaCl, 1 mM MgCl_2_, 5% glycerol, 0.15 mM phenylmethylsulfonyl fluoride (PMSF), 0.5 mM Tris(2-carboxyethyl)phosphine (TCEP), and protease inhibitor cocktail (Roche). Resuspended cells were lysed by sonication on ice, and lysate was precleared by centrifugation for 30 min at 4 °C at 25,000g. MBP-tagged BC was purified via amylose affinity chromatography in a gravity flow column setup. The lysate supernatant was applied to pre-equilibrated resin and allowed to batch bind at 4°C with rotation. The resin was washed with 50 mM Tris pH 7.5, 150 mM NaCl, 1 mM MgCl_2_, 5% glycerol, 0.5 mM TCEP. Elution was performed with 10 mM maltose dissolved in wash buffer. Subsequently, TEV protease was added to purified MBP-tev-BC within dialysis tubing. Sample was incubated overnight at 4°C. The TEV-cleaved BC sample was separated via size exclusion chromatography in 100 mM HEPES pH 8, 4 mM MgCl_2_, 0.5 mM TCEP on a Superdex 200 Increase 10/300 column (Cytiva).

### 4.2. Cryo-EM

Purified BC was concentrated to 1.2 mg mL^-1^ and 3.5 µL of sample was applied onto glow discharged UltrAuFoil R1.2/1.3 300 mesh grids using a VitroBot Mark IV Plus (Thermo Fisher Scientific). Sample was applied at 4°C, 100% humidity. Grids were blotted for 3 s with blot force 12 and plunge frozen into liquid ethane. Data was collected on a Titan Krios G4 transmission electron microscope (TEM) operating at 300 kV, equipped with a Falcon 4 Direct Electron Detector. Frames were collected at 130kX magnification and a pixel size of 0.97 Å/pix with a target defocus range of-1.6 µm to-0.6 µm and a total exposure of 40e^-^/ Å^2^. In total, 5520 micrographs were collected, which were imported into cryoSPARC v4.6.2.

Movies were motion corrected using the Patch Motion Correction program, and calculated by Patch CTF. Initial particle picking was done with Blob Picker with a diameter range of 120 to 160 Å. These particles were used to generate 2D classes, and selected 2D classes were used for template-based particle selection. A total of 3,455, 841 particles were extracted with a box size of 256 and were refined over several rounds of 2D classification. After 2D classification, 142,226 high quality particles were used for Ab-initio reconstitution. Global CTF Refinement was performed on the particles to adjust higher-order CTF terms, and then non-uniform refinement was applied with C2 symmetry. The final map of the BC dimer was refined to 2.85 Å using the gold-standard FSC=0.143 standard.

For model building, the AlphaFold3 predicted model of the BC dimer was docked into the cryo-EM map using ChimeraX, and then subsequently refined in real space using Coot, ISOLDE, and Phenix iteratively. All structural models shown were prepared using PyMol and ChimeraX. The cryo-EM tetramer map was of insufficient resolution for modeling; two copies of the BC dimer model fit into the tetramer cryo-EM map.

### 4.3. Ranking the importance of BC residues

The Evolutionary Trace (ET) method of analyzing a multiple sequence alignment (MSA) alignment successfully predicts functional surfaces of a protein (59). A hybrid method known as real-valued ET (rvET) combines quantitative scoring by ET and information entropy of the MSA. Because rvET outperformed ET and entropy and was more robust to flaws in the MSA (60), we used rvET to predict functional surfaces of BC. We submitted to the rvET server (61) the sequence of BC from pennycress, Arabidopsis, or the cryo-EM structure of pennycress BC. The server performed a BLAST search against the UniProt UniRef90 library of sequences low in redundancy. This provided significant alignments to 500 unique homologues (listed in Table S2). The rvET server used the MSA to rank evolutionary importance of each position in sequence (Table S3, Fig. S5).

### 4.4. Comparative structural bioinformatics

The structural bioinformatics routines of Bio3D (39) were used as a package in R (version 4.1.2) operated using the RStudio environment. Bio3D identified and aligned homologous structures of BC using its *blast.pdb*, *pdbaln*, *seqidentity*, and *rmsd* functions. The experimental structures of BC subunits most like the cryo-EM structure of pennycress BC proved to be from bacteria, which included the thermophiles *C. auranticus*, *H. thermophilus*, and *A. aeolicus*. The structures varied in the active site being vacant or occupied by bicarbonate and biotin substrates, non-hydrolyzable ATP analogue, or ADP.

The *core* function was used to find the core of residues shared among the 11 BC enzymes analyzed. The domains and sub-domains that move as a unit within the BC enzymes were identified using the GeoStaS algorithm (40) implemented in Bio3D (39). These domains were identified across the shared, core sequence by the plot of high atomic movement similarity matrix.

Principal component analysis of the atomic structural models using the *pca* function of Bio3D, followed by hierarchical clustering using the *hclust* function, suggested the similarity of the models (39). Normal modes analysis of the ensemble of 11 structures used the *nma* function which by default uses the Cα forcefield based on Amber94 (39, 45). This jointly identified the most significant modes of motion and enabled comparison of the motions across the ensemble (39). Movies depicting the top normal modes were constructed using the *mktrj.enma* and *geostas* functions to generate series of backbone Cα traces. Movies were made from these frames using Pymol.

### 4.5. Surface accessibility by acetylation, i.e. protein painting

The acetylation reactions followed Barth et al. (46). Stock solutions of NHS-Acetate were prepared at 10 mM in 25% acetonitrile and 2 mM in 5% acetonitrile. For each reaction, the protein solution at a final concentration of BC protomers of 3 to 10 μM was incubated with the acetylation reagent in 40 μL at 23°C for 40 min. The reactions were then quenched with 2 μL of 100 mM Tris-HCl and frozen at-20°C. The concentrations of NHS-acetate used in the acetylation reactions were 0, 70, 200, 600, 1800 μM. These molar excesses of 7, 20, 60, and 180-fold over the 10 μM BC samples correspond to 0.3, 0.9, 2.7, and 8.2-fold the 22 Lysine residues per chain of BC. Acetylation reactions were performed using one to four biological replicates.

The acetylated peptide fragments were isolated and detected in a typical proteomic workflow (40). The acetylated BC was reduced with 10 mM DTT in 10 mM ammonium bicarbonate at 30°C for 30 min and then alkylated with 40 mM iodoacetamide in 10 mM ammonium bicarbonate for 1 h at room temperature. Glu-C (Promega) digestion was initiated by adding the enzyme at a 1:12 ratio to protein and incubating the samples at 37°C for 14 hours. To improve digestion efficiency, an additional dose of Glu-C was then added, followed by another 6-hour incubation, bringing the Glu-C to the acetylated, reduced, and alkylated BC to a 1:6 final ratio. The peptide digest was acidified to a final 0.1% formic acid (v/v), submitted to desalting step with Pierce® C18 Tips, lyophilized using a centrifugal evaporator, and stored at-80°C until mass spectrometry. The lyophilized samples were resuspended in 32 μL of 0.1% (v/v) formic acid in water and two different workflows were performed for data acquisition and database search steps.

Firstly, data acquisition parameters were optimized to maximize peptide coverage and ensure reproducibility across injections. Each sample was first loaded onto an OPTI-TRAP™ cartridge (Optimize Technologies, Oregon City, OR; 5 µL, 0.5 mm × 1.3 mm) and subsequently eluted onto a self-packed analytical column (20 cm × 75 µm internal diameter, HxSil 5-µm C18 matrix). Peptides were separated using a Finnigan Surveyor liquid chromatography (LC) system operating at 125 μL/min in a split-flow configuration over a 60-min gradient. The mobile phases consisted of 0.1% formic acid in water (phase A) and 0.1% formic acid in acetonitrile (phase B). Data were acquired on an LTQ Orbitrap XL mass spectrometer (Thermo Fisher, San Jose, CA) in data-dependent acquisition (DDA) mode, selecting the top 11 most abundant precursor ions from each MS1 scan for fragmentation. MS1 spectra were acquired in positive-ion FTMS mode at 60,000 resolution over an m/z range of 200–2000 using centroid data. Peptides were fragmented by collision-induced dissociation (CID) with a collision energy of 35 kV, an isolation width of 2.0 m/z, an activation time of 30 ms, and a minimum signal threshold of 500. Only charge states ≥+2 were selected for fragmentation. Dynamic exclusion was enabled with a repeat count of 1, a repeat duration of 30 s, an exclusion list size of 100, and an exclusion duration of 30 s. Raw files were searched individually in Proteome Discoverer 2.4 against the pennycress protein database (26,393 entries, BioProject PRJEB46635) containing BC (CAH2072050.1) protein sequence. Spectra were filtered based on MS1 precursor masses between 350 and 5000 Da. High-confidence identifications were obtained using the Sequest HT search algorithm with a precursor mass tolerance of 5 ppm and a fragment mass tolerance of 0.6 Da. Up to five dynamic modifications were permitted per peptide, including methionine oxidation (+15.995 Da) and acetylation of lysine, serine, threonine, and tyrosine residues (+42.011 Da). Validation of protein and peptide-spectrum matches (PSMs) was performed with a Target–Decoy approach, applying an FDR threshold of ≤0.01 for high-confidence identifications and ≤0.05 for medium-confidence identifications. Phosphorylation site localization was refined using the IMP-ptmRS tool.

Secondly, peptide separation was performed using an EvoSep ultrahigh-performance liquid chromatography system with reverse-phase chromatography over a 44-min gradient at a flow rate of 100 nL/min. Peptides were resolved on a C18 analytical column (PepSep C18, Bruker Daltonics; 15 cm × 75 μm, 1.9-μm particle size). Mass spectra were acquired online during chromatographic separation on a TimsTOF Pro 2 (Bruker Daltonics) operated in positive-ion mode using a data-dependent PASEF method, with an m/z acquisition range of 100–1700. Both PASEF and TIMS functionalities were enabled. One mass spectrum and ten PASEF frames were obtained per cycle of 1.17sec. The target MS/MS intensity was set to 10,000 with a minimum threshold of 2,500. Collision energies ranged from 20 to 59 eV and were applied using a charge-state–dependent rolling collision energy scheme. An active precursor exclusion and reconsideration strategy was used, with release after 0.4 min, and a second MS/MS event was triggered when precursor intensity increased at least four-fold within a mass tolerance of 0.015 m/z. Automated *de novo* sequencing of MS/MS spectra was carried out using PEAKS Studio 10.0 (Bioinformatics Solutions, Toronto, Canada), with precursor and fragment mass error tolerances set to 20 ppm and 0.05 Da, respectively. Variable modifications were included as described previously. PSM identification was subsequently performed with the PEAKS DB module, using the built-in target–decoy strategy to maintain a 1% false discovery rate. Phosphorylation site localization confidence was assessed using the Ascore algorithm, applying a minimum threshold of 20. The mass spectrometry proteomics data have been deposited to the ProteomeXchange Consortium via the PRIDE partner repository (47) with the dataset identifier PXD071146.

The proportion of each sidechain that was acetylated (mainly lysine sidechains) was calculated as described (46):

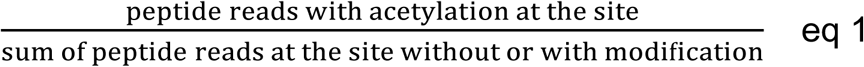

To measure this ratio at each site, we wrote a Python script, ResiCritic, which aligned each peptide sequence to the protein FASTA sequence and counted how many times each residue was found acetylated and found non-acetylated. The script created a coverage file and a modification file listing the number of times the residue was found modified, the number of times this residue was found without or with modification, and their ratio specified in eq. 1. ResiCritic is available at doi.org/10.5281/zenodo.18040770

### 4.6. Cross-linking mass spectrometry to probe oligomers

The cross-linking reactions leveraged cleavage of the DSBU (disuccinimidyl dibutyric urea) crosslinker during the collision induced dissociation (CID) of MS-MS measurements (48–53). This generated fragment ions and patterns for automated recognition of cross-links. The Tris buffer of BC was exchanged for 50 mM HEPES, 150 mM NaCl, pH 7.5, using a Micro Bio-Spin chromatography column (Bio-Rad).BC (0.7 mg/ml) was incubated with 1 mM DSBU for 1 h at room temperature to introduce cross-links. The reactions were quenched by adding Tris to a final concentration of 20 mM. Cross-linked BC samples were separated by SDS-PAGE, and BC bands at approximately 100 kDa and 250 kDa were reduced with DTT and destained with 100 mM ammonium bicarbonate/acetonitrile (1:1, v/v). Proteins in the bands were digested with trypsin overnight at 37 °C with agitation at 400 rpm on a Thermomixer; the digestion volume was sufficient to cover the gel pieces. Peptides were extracted using 5% formic acid/acetonitrile (1:2, v/v) by incubating for 15 min at 37 °C with shaking. Cross-linked BC samples were also directly denatured with 8 M urea. After denaturation, proteins were reduced with 5 mM DTT for 1 h at 25 °C, followed by alkylation with 40 mM iodoacetamide for 45 min at 25 °C in the dark. Samples were diluted three-fold with 50 mM ammonium bicarbonate and digested with trypsin at an enzyme:substrate ratio of 1:50 (w/w) for 16 h at 25 °C with shaking at 850 rpm. Protease activity was quenched by adding 0.1% formic acid. Digested peptides were applied to microspin C18 cartridges (NEST Group), desalted, and eluted with 80% acetonitrile, 0.1% TFA. Eluates were dried and stored at −20 °C until LC-MS/MS analysis.

For mass spectrometry data acquisition, cross-linked peptides were analyzed on a Q Exactive HF-X Orbitrap mass spectrometer (Thermo Fisher Scientific). Peptides were reconstituted in 0.1% formic acid and separated on a Thermo Vanquish Neo LC with an in-house packed Waters AQ C18 analytical column (30 cm × 75 µm i.d., 1.7 µm). Chromatography was performed at 0.2 µL/min using mobile phase A (0.1% formic acid in H2O) and mobile phase B (0.1% formic acid in acetonitrile). Peptides were eluted over 161 min using the following gradients (time in minutes, %B): 0–4, 1% B; 4–14, 1–8% B; 14–108, 8–21% B; 108–133, 21–28% B; 133–145, 28–37% B; 145–151, 37–75% B; 151–161, 75–95% B. MS1 spectra were acquired from 300–1800 m/z at a resolution of 60K. MS/MS spectra were acquired in data-dependent mode using a 0.7 m/z isolation window at a resolution of 45K. Peptides were fragmented by HCD at a normalized collision energy of 30, with a 45 s dynamic exclusion, analyzing charge states +2 to +6.

Cross-link spectrum matches (CSMs) were identified using pLink (version 3.0.16) (54, 55) and Scout (version 1.6.2) (56). Searches were performed against custom FASTA containing sequences for *Thlaspi arvense* BC, BADC1, and BCCP with the false discovery rate (FDR) set to 5%.

### 4.7. Structural modeling guided by cross-links

Elucidation of the oligomeric structure of a homo-oligomer is complicated by the ambiguity of identifying crosslinks between chains. We addressed this challenge by dividing it into two phases of iterative structural interpretation. In both phases, we sought to identify sets of interchain crosslinks that were compatible with one another and a structural model. The first phase was simplified by structural modeling using only two of the four chains of BC in a tetramer, i.e. two protomers from separate dimers. Distance restraints corresponding to varied combinations of potentially crosslinked residues were incorporated into High Ambiguity Driven Docking calculations on the HADDOCK 2.4 server (43). This comprised HADDOCK’s standard stages of rigid body docking from randomized orientations, followed by torsion angle dynamics with semi-flexibility, and then refinement in water. The HADDOCK server operates in the cloud in the WeNMR-EOSC Ecosystem (57). After the first phase identified four mutually compatible interchain crosslinks (Fig. 4A), the second phase expanded the two protomers that had been docked with the core crosslinks, into two dimers. The two additional chains suggested more areas of potential proximity. The four remaining candidates for interchain crosslinks were tested for compatibility with the first four interchain crosslinks identified using HADDOCK calculations. The lysine residues found to be engaged in probable cross-links between chains were allowed full-flexibility in the final structural refinement using the seven crosslinked-distance restraints. The rotatable lid domain (B-domain, residues 200 - 275) in contact with another protomer were allowed semi-flexibility in the final refinement. Of 600 models (structural variants) calculated initially, 120 were selected for refinement with flexibility and in a water environment. Seven of these were chosen as the final set of structural variants, because they were free of violations of the crosslink-based distance restraints (but for a 0.8 Å violation in one case). Moreover, the seven were also selected to be in the upper quartile of 120 in HADDOCK score and upper half of the 120 in sum of all stereochemical energy terms, including non-bonded van der Waals and electrostatic interactions. The ensemble of seven comprised two clusters of slightly different binding poses (Table 2) which we compared quantitatively using ChimeraX (58).

## 5. Data Availability

All primary data will be deposited in online databases. The Cryo Electron Microscopy maps and structure models will be deposited in the EMDB and PDB, respectively (EMDB: 74880, 74881 and PDB: 9ZVM). The structural models of the cross-linked dimer-of-dimers will be deposited at PDB-IHM (PDB-IHM: XXXXX, PDB: XXXXX). The proteomics raw data of the acetylation study are deposited in PRIDE under accession code PXD071146. The cross-linking data will be deposited as well. The script (ResiCritic) used to quantify the acetylation is available at GitHub and Zenodo (doi.org/10.5281/zenodo.18040770).

## Acknowledgments

This work received support from Environmental Molecular Sciences Laboratory project 61285 (SRV, JJT), a Univ. of Missouri College of Agriculture Joy of Discovery award (SRV, ALY), DOE grant DE-SC0023142 (JJT), and Hatch Multi-State Project No. 1013013 from the National Institute of Food and Agriculture (SRV, JJT, ALY), and Univ. of University of Missouri startup funds (ALY). Initially cryo-electron microscopy screening was supported by the Kansas University Medical Center Cryo-electron microscopy Facility and the Glacios equipment purchase via NIH grant S10 OD036339-01.

## Literature cited

1. Thelen, J. J., and Ohlrogge, J. B. (2002) Metabolic Engineering of Fatty Acid Biosynthesis in Plants. Metabolic Engineering. 4, 12–21

2. Raboanatahiry, N., Li, H., Yu, L., and Li, M. (2021) Rapeseed (Brassica napus): Processing, Utilization, and Genetic Improvement. Agronomy. 11, 1776

3. So, K. K. Y., and Duncan, R. W. (2021) Breeding Canola (Brassica napus L.) for Protein in Feed and Food. Plants. 10, 2220

4. Nunn, A., Rodríguez-Arévalo, I., Tandukar, Z., Frels, K., Contreras-Garrido, A., Carbonell-Bejerano, P., Zhang, P., Ramos Cruz, D., Jandrasits, K., Lanz, C., Brusa, A., Mirouze, M., Dorn, K., Galbraith, D. W., Jarvis, B. A., Sedbrook, J. C., Wyse, D. L., Otto, C., Langenberger, D., Stadler, P. F., Weigel, D., Marks, M. D., Anderson, J. A., Becker, C., and Chopra, R. (2022) Chromosome-level *Thlaspi arvense* genome provides new tools for translational research and for a newly domesticated cash cover crop of the cooler climates. Plant Biotechnology Journal. 20, 944–963

5. Guchhait, R. B., Polakis, S. E., Dimroth, P., Stoll, E., Moss, J., and Lane, M. D. (1974) Acetyl Coenzyme A Carboxylase System of Escherichia coli. Journal of Biological Chemistry. 249, 6633–6645

6. Cronan, J. E., and Waldrop, G. L. (2002) Multi-subunit acetyl-CoA carboxylases. Progress in Lipid Research. 41, 407–435

7. Konishi, T., and Sasaki, Y. (1994) Compartmentalization of two forms of acetyl-CoA carboxylase in plants and the origin of their tolerance toward herbicides. Proc. Natl. Acad. Sci. U.S.A. 91, 3598–3601

8. Konishi, T., Shinohara, K., Yamada, K., and Sasaki, Y. (1996) Acetyl-CoA Carboxylase in Higher Plants: Most Plants Other Than Gramineae Have Both the Prokaryotic and the Eukaryotic Forms of This Enzyme. Plant and Cell Physiology. 37, 117–122

9. Reverdatto, S., Beilinson, V., and Nielsen, N. C. (1999) A Multisubunit Acetyl Coenzyme A Carboxylase from Soybean1. Plant Physiology. 119, 961–978

10. Chou, C.-Y., Yu, L. P. C., and Tong, L. (2009) Crystal Structure of Biotin Carboxylase in Complex with Substrates and Implications for Its Catalytic Mechanism. Journal of Biological Chemistry. 284, 11690–11697

11. Perham, R. N. (2000) Swinging Arms and Swinging Domains in Multifunctional Enzymes: Catalytic Machines for Multistep Reactions. Annu. Rev. Biochem. 69, 961–1004

12. Salie, M. J., Zhang, N., Lancikova, V., Xu, D., and Thelen, J. J. (2016) A Family of Negative Regulators Targets the Committed Step of de Novo Fatty Acid Biosynthesis. Plant Cell. 28, 2312–2325

13. Salie, M. J., and Thelen, J. J. (2016) Regulation and structure of the heteromeric acetyl-CoA carboxylase. Biochimica et Biophysica Acta (BBA) - Molecular and Cell Biology of Lipids. 1861, 1207–1213

14. Conrado, A. C., Lemes Jorge, G., Rao, R. S. P., Xu, C., Xu, D., Li-Beisson, Y., and Thelen, J. J. (2024) Evolution of the regulatory subunits for the heteromeric acetyl-CoA carboxylase. Phil. Trans. R. Soc. B. 10.1098/rstb.2023.0353

15. Ye, Y., Fulcher, Y. G., Sliman, D. J., Day, M. T., Schroeder, M. J., Koppisetti, R. K., Bates, P. D., Thelen, J. J., and Van Doren, S. R. (2020) The BADC and BCCP subunits of chloroplast acetyl-CoA carboxylase sense the pH changes of the light–dark cycle. Journal of Biological Chemistry. 295, 9901–9916

16. Garneau, M. G., Jorge, G. L., Shockey, J., Thelen, J. J., and Bates, P. D. (2024) PII interactions with BADC and BCCP proteins co-regulate lipid and nitrogen metabolism in Arabidopsis. 10.1101/2024.11.04.621944

17. Waldrop, G. L., Rayment, I., and Holden, H. M. (1994) Three-Dimensional Structure of the Biotin Carboxylase Subunit of Acetyl-CoA Carboxylase. Biochemistry. 33, 10249–10256

18. Thoden, J. B., Blanchard, C. Z., Holden, H. M., and Waldrop, G. L. (2000) Movement of the Biotin Carboxylase B-domain as a Result of ATP Binding. Journal of Biological Chemistry. 275, 16183–16190

19. Shen, Y., Chou, C.-Y., Chang, G.-G., and Tong, L. (2006) Is Dimerization Required for the Catalytic Activity of Bacterial Biotin Carboxylase? Molecular Cell. 22, 807–818

20. Cheng, C. C., Shipps, G. W., Yang, Z., Sun, B., Kawahata, N., Soucy, K. A., Soriano, A., Orth, P., Xiao, L., Mann, P., and Black, T. (2009) Discovery and optimization of antibacterial AccC inhibitors. Bioorganic & Medicinal Chemistry Letters. 19, 6507–6514

21. Chou, C.-Y., and Tong, L. (2011) Structural and Biochemical Studies on the Regulation of Biotin Carboxylase by Substrate Inhibition and Dimerization. Journal of Biological Chemistry. 286, 24417–24425

22. Kondo, S., Nakajima, Y., Sugio, S., Yong-Biao, J., Sueda, S., and Kondo, H. (2004) Structure of the biotin carboxylase subunit of pyruvate carboxylase from *Aquifex aeolicus* at 2.2 Å resolution. Acta Crystallogr D Biol Crystallogr. 60, 486–492

23. Buhrman, G., Enríquez, P., Dillard, L., Baer, H., Truong, V., Grunden, A. M., and Rose, R. B. (2021) Structure, Function, and Thermal Adaptation of the Biotin Carboxylase Domain Dimer from *Hydrogenobacter thermophilus* 2-Oxoglutarate Carboxylase. Biochemistry. 60, 324–345

24. Mochalkin, I., Miller, J. R., Evdokimov, A., Lightle, S., Yan, C., Stover, C. K., and Waldrop, G. L. (2008) Structural evidence for substrate-induced synergism and half-sites reactivity in biotin carboxylase. Protein Science. 17, 1706–1718

25. Broussard, T. C., Pakhomova, S., Neau, D. B., Bonnot, R., and Waldrop, G. L. (2015) Structural Analysis of Substrate, Reaction Intermediate, and Product Binding in *Haemophilus influenzae* Biotin Carboxylase. Biochemistry. 54, 3860–3870

26. Andrews, L. D., Kane, T. R., Dozzo, P., Haglund, C. M., Hilderbrandt, D. J., Linsell, M. S., Machajewski, T., McEnroe, G., Serio, A. W., Wlasichuk, K. B., Neau, D. B., Pakhomova, S., Waldrop, G. L., Sharp, M., Pogliano, J., Cirz, R. T., and Cohen, F. (2019) Optimization and Mechanistic Characterization of Pyridopyrimidine Inhibitors of Bacterial Biotin Carboxylase. J. Med. Chem. 62, 7489–7505

27. Mavila, A. M., Vargas, J. A., Condori, E., Suclupe Farro, E. G., Furtado, A. A., López, J. M., Gonzalez, S. L., Pereira, H. D., Marapara, J. L., Paredes, R. R., Cobos, M., Castro, J. C., Garratt, R. C., and Leonardo, D. A. (2025) Phylogenetic analysis and structural studies of heteromeric acetyl-CoA carboxylase from the oleaginous Amazonian microalgae Ankistrodesmus sp.: Insights into the BC and BCCP subunits. Journal of Structural Biology. 217, 108200

28. Lee, J. K. J., Liu, Y.-T., Hu, J. J., Aphasizheva, I., Aphasizhev, R., and Zhou, Z. H. (2023) CryoEM reveals oligomeric isomers of a multienzyme complex and assembly mechanics. Journal of Structural Biology: X. 7, 100088

29. Shivaiah, K.-K., Ding, G., Upton, B., and Nikolau, B. J. (2020) Non-Catalytic Subunits Facilitate Quaternary Organization of Plastidic Acetyl-CoA Carboxylase. Plant Physiol. 182, 756–775

30. Xu, X., Silva De Sousa, A., Boram, T. J., Jiang, W., and Lohman, J. R. (2024) ActiveE. coliheteromeric acetyl-CoA carboxylase forms polymorphic helical tubular filaments. 10.1101/2024.05.28.596234

31. Thoden, J. B., Blanchard, C. Z., Holden, H. M., and Waldrop, G. L. (2000) Movement of the Biotin Carboxylase B-domain as a Result of ATP Binding. Journal of Biological Chemistry. 275, 16183–16190

32. Shen, Y., Chou, C.-Y., Chang, G.-G., and Tong, L. (2006) Is Dimerization Required for the Catalytic Activity of Bacterial Biotin Carboxylase? Molecular Cell. 22, 807–818

33. Chou, C.-Y., Yu, L. P. C., and Tong, L. (2009) Crystal Structure of Biotin Carboxylase in Complex with Substrates and Implications for Its Catalytic Mechanism. Journal of Biological Chemistry. 284, 11690–11697

34. Broussard, T. C., Kobe, M. J., Pakhomova, S., Neau, D. B., Price, A. E., Champion, T. S., and Waldrop, G. L. (2013) The Three-Dimensional Structure of the Biotin Carboxylase-Biotin Carboxyl Carrier Protein Complex of E. coli Acetyl-CoA Carboxylase. Structure. 21, 650–657

35. Andrews, L. D., Kane, T. R., Dozzo, P., Haglund, C. M., Hilderbrandt, D. J., Linsell, M. S., Machajewski, T., McEnroe, G., Serio, A. W., Wlasichuk, K. B., Neau, D. B., Pakhomova, S., Waldrop, G. L., Sharp, M., Pogliano, J., Cirz, R. T., and Cohen, F. (2019) Optimization and Mechanistic Characterization of Pyridopyrimidine Inhibitors of Bacterial Biotin Carboxylase. J. Med. Chem. 62, 7489–7505

36. Waldrop, G. L., Rayment, I., and Holden, H. M. Three-Dimensional Structure of the Biotin Carboxylase Subunit of Acetyl-CoA Carboxylase

37. Broussard, T. C., Kobe, M. J., Pakhomova, S., Neau, D. B., Price, A. E., Champion, T. S., and Waldrop, G. L. (2013) The Three-Dimensional Structure of the Biotin Carboxylase-Biotin Carboxyl Carrier Protein Complex of E. coli Acetyl-CoA Carboxylase. Structure. 21, 650–657

38. Shen, J., Wu, W., Wang, K., Wu, J., Liu, B., Li, C., Gong, Z., Hong, X., Fang, H., Zhang, X., and Xu, X. (2024) *Chloroflexus aurantiacus* acetyl-CoA carboxylase evolves fused biotin carboxylase and biotin carboxyl carrier protein to complete carboxylation activity. mBio. 15, e03414–23

39. Skjærven, L., Yao, X.-Q., Scarabelli, G., and Grant, B. J. (2014) Integrating protein structural dynamics and evolutionary analysis with Bio3D. BMC Bioinformatics. 15, 399

40. Romanowska, J., Nowiński, K. S., and Trylska, J. (2012) Determining Geometrically Stable Domains in Molecular Conformation Sets. J. Chem. Theory Comput. 8, 2588–2599

41. Novak, P., Kruppa, G. H., Young, M. M., and Schoeniger, J. (2004) A Top-down method for the determination of residue-specific solvent accessibility in proteins. J. Mass Spectrom. 39, 322–328

42. Barth, M., Bender, J., Kundlacz, T., and Schmidt, C. (2020) Evaluation of NHS-Acetate and DEPC labelling for determination of solvent accessible amino acid residues in protein complexes. Journal of Proteomics. 222, 103793

43. Honorato, R. V., Trellet, M. E., Jiménez-García, B., Schaarschmidt, J. J., Giulini, M., Reys, V., Koukos, P. I., Rodrigues, J. P. G. L. M., Karaca, E., Van Zundert, G. C. P., Roel-Touris, J., Van Noort, C. W., Jandová, Z., Melquiond, A. S. J., and Bonvin, A. M. J. J. (2024) The HADDOCK2.4 web server for integrative modeling of biomolecular complexes. Nat Protoc. 19, 3219–3241

44. Almagro Armenteros, J. J., Salvatore, M., Emanuelsson, O., Winther, O., Von Heijne, G., Elofsson, A., and Nielsen, H. (2019) Detecting sequence signals in targeting peptides using deep learning. Life Sci. Alliance. 2, e201900429

45. Hinsen, K., Petrescu, A.-J., Dellerue, S., Bellissent-Funel, M.-C., and Kneller, G. R. (2000) Harmonicity in slow protein dynamics. Chemical Physics. 261, 25–37

46. Barth, M., Bender, J., Kundlacz, T., and Schmidt, C. (2020) Evaluation of NHS-Acetate and DEPC labelling for determination of solvent accessible amino acid residues in protein complexes. Journal of Proteomics. 222, 103793

47. Perez-Riverol, Y., Bandla, C., Kundu, D. J., Kamatchinathan, S., Bai, J., Hewapathirana, S., John, N. S., Prakash, A., Walzer, M., Wang, S., and Vizcaíno, J. A. (2025) The PRIDE database at 20 years: 2025 update. Nucleic Acids Research. 53, D543–D553

48. Müller, M. Q., Zeiser, J. J., Dreiocker, F., Pich, A., Schäfer, M., and Sinz, A. (2011) A universal matrix-assisted laser desorption/ionization cleavable cross-linker for protein structure analysis. Rapid Comm Mass Spectrometry. 25, 155–161

49. Iacobucci, C., Götze, M., Ihling, C. H., Piotrowski, C., Arlt, C., Schäfer, M., Hage, C., Schmidt, R., and Sinz, A. (2018) A cross-linking/mass spectrometry workflow based on MS-cleavable cross-linkers and the MeroX software for studying protein structures and protein–protein interactions. Nat Protoc. 13, 2864–2889

50. Müller, M. Q., Dreiocker, F., Ihling, C. H., Schäfer, M., and Sinz, A. (2010) Cleavable Cross-Linker for Protein Structure Analysis: Reliable Identification of Cross-Linking Products by Tandem MS. Anal. Chem. 82, 6958–6968

51. Hage, C., Iacobucci, C., Rehkamp, A., Arlt, C., and Sinz, A. (2017) The First Zero-Length Mass Spectrometry-Cleavable Cross-Linker for Protein Structure Analysis. Angew Chem Int Ed. 56, 14551–14555

52. Arlt, C., Götze, M., Ihling, C. H., Hage, C., Schäfer, M., and Sinz, A. (2016) Integrated Workflow for Structural Proteomics Studies Based on Cross-Linking/Mass Spectrometry with an MS/MS Cleavable Cross-Linker. Anal. Chem. 88, 7930–7937

53. Piersimoni, L., Kastritis, P. L., Arlt, C., and Sinz, A. (2022) Cross-Linking Mass Spectrometry for Investigating Protein Conformations and Protein–Protein Interactions─A Method for All Seasons. Chem. Rev. 122, 7500–7531

54. Fan, S., Meng, J., Lu, S., Zhang, K., Yang, H., Chi, H., Sun, R., Dong, M., and He, S. (2015) Using pLink to Analyze Cross-Linked Peptides. CP in Bioinformatics. 10.1002/0471250953.bi0821s49

55. Barysz, H. M., and Malmström, J. (2018) Development of Large-scale Cross-linking Mass Spectrometry. Molecular & Cellular Proteomics. 17, 1055–1066

56. Clasen, M. A., Ruwolt, M., Wang, C., Ruta, J., Bogdanow, B., Kurt, L. U., Zhang, Z., Wang, S., Gozzo, F. C., Chen, T., Carvalho, P. C., Lima, D. B., and Liu, F. (2024) Proteome-scale recombinant standards and a robust high-speed search engine to advance cross-linking MS-based interactomics. Nat Methods. 21, 2327–2335

57. Honorato, R. V., Koukos, P. I., Jiménez-García, B., Tsaregorodtsev, A., Verlato, M., Giachetti, A., Rosato, A., and Bonvin, A. M. J. J. (2021) Structural Biology in the Clouds: The WeNMR-EOSC Ecosystem. Front. Mol. Biosci. 8, 729513

58. Pettersen, E. F., Goddard, T. D., Huang, C. C., Meng, E. C., Couch, G. S., Croll, T. I., Morris, J. H., and Ferrin, T. E. (2021) UCSF CHIMERAX: Structure visualization for researchers, educators, and developers. Protein Science. 30, 70–82

59. Lichtarge, O., Bourne, H. R., and Cohen, F. E. (1996) An Evolutionary Trace Method Defines Binding Surfaces Common to Protein Families. Journal of Molecular Biology. 257, 342–358

60. Mihalek, I., Reš, I., and Lichtarge, O. (2004) A Family of Evolution–Entropy Hybrid Methods for Ranking Protein Residues by Importance. Journal of Molecular Biology. 336, 1265–1282

61. Wilkins, A., Erdin, S., Lua, R., and Lichtarge, O. (2012) Evolutionary Trace for Prediction and Redesign of Protein Functional Sites. in Computational Drug Discovery and Design (Baron, R. ed), pp. 29–42, Methods in Molecular Biology, Springer New York, New York, NY, 819, 29–42

62. Grant, B. J., Rodrigues, A. P. C., ElSawy, K. M., McCammon, J. A., and Caves, L. S. D. (2006) Bio3d: an R package for the comparative analysis of protein structures. Bioinformatics. 22, 2695–2696

63. Skjærven, L., Yao, X.-Q., Scarabelli, G., and Grant, B. J. (2014) Integrating protein structural dynamics and evolutionary analysis with Bio3D. BMC Bioinformatics. 15, 399

