## Supplemental Figures for "Oligomeric assemblies of plant biotin carboxylase revealed by cryo-EM and cross-linking"

### Supporting Information list

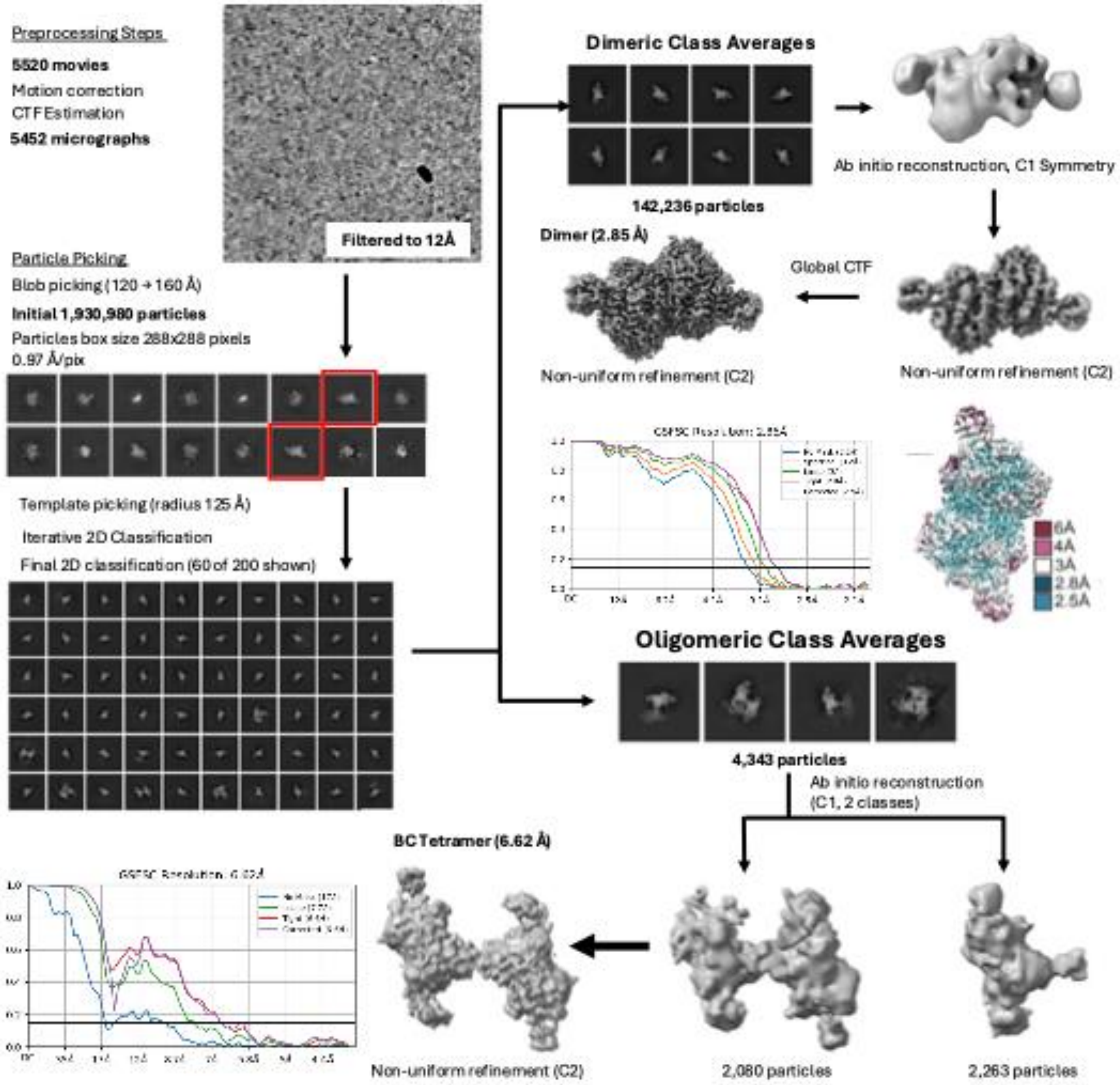


Figure S1. Cryo-EM workflow with intermediate results for pennycress BC, including particle counts, 2D class averages, refinement steps, and map quality and resolution.

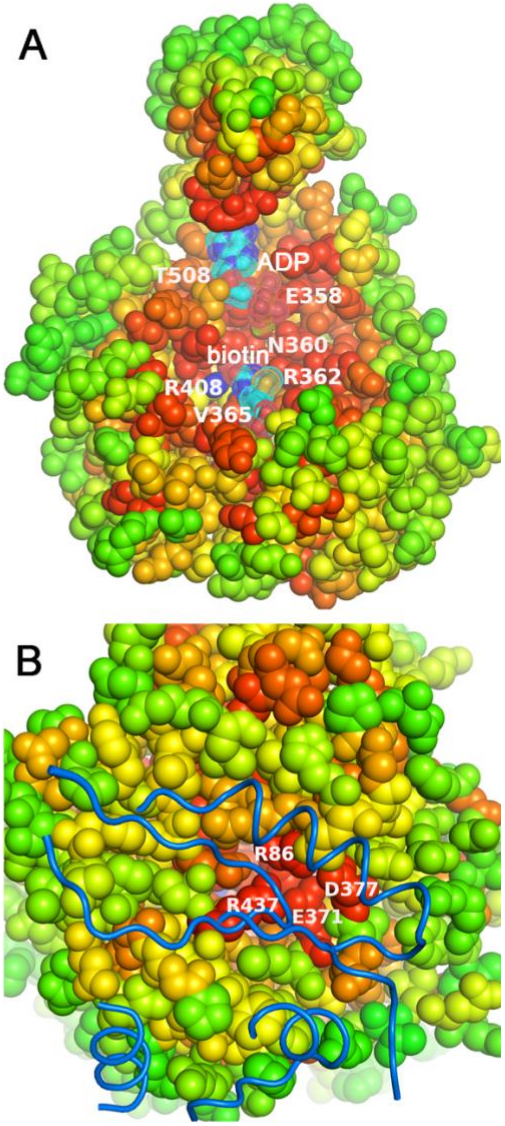


Figure S2. Conservation among 500 BC catalytic subunits mapped onto (A) the active site and (B) the dimer interface. The conservation was scored by real-valued Evolutionary Trace (rvET) analysis. Details of the rvET are provided in Methods, Table S3, and Fig. S6. Red represents the highest conservation and green the lowest conservation.

A

| | 3RUP | 3RV4 | 2VPQ | 7KCT | 4MV1 | 7KBL | TABC | 2C00 | 1ULZ | 1BNC | 8HZ4 |
|---|---|---|---|---|---|---|---|---|---|---|---|
| **3RUP** | 1.000 | 0.998 | 0.534 | 0.530 | 0.814 | 0.528 | **0.537** | 0.712 | 0.529 | 1.000 | 0.535 |
| **3RV4** | 0.998 | 1.000 | 0.530 | 0.527 | 0.812 | 0.523 | **0.535** | 0.711 | 0.527 | 0.998 | 0.541 |
| **2VPQ** | 0.534 | 0.530 | 1.000 | 0.552 | 0.546 | 0.549 | **0.538** | 0.515 | 0.561 | 0.532 | 0.502 |
| **7KCT** | 0.530 | 0.527 | 0.552 | 1.000 | 0.530 | 0.999 | **0.545** | 0.509 | 0.825 | 0.526 | 0.503 |
| **4MV1** | 0.814 | 0.812 | 0.546 | 0.530 | 1.000 | 0.531 | **0.526** | 0.713 | 0.523 | 0.813 | 0.550 |
| **7KBL** | 0.528 | 0.523 | 0.549 | 0.999 | 0.531 | 1.000 | **0.539** | 0.507 | 0.822 | 0.525 | 0.503 |
| **TABC** | **0.537** | **0.535** | **0.538** | **0.545** | **0.526** | **0.539** | **1.000** | **0.518** | **0.534** | **0.531** | **0.524** |
| **2C00** | 0.712 | 0.711 | 0.515 | 0.509 | 0.713 | 0.507 | **0.518** | 1.000 | 0.523 | 0.715 | 0.509 |
| **1ULZ** | 0.529 | 0.527 | 0.561 | 0.825 | 0.523 | 0.822 | **0.534** | 0.523 | 1.000 | 0.529 | 0.510 |
| **1BNC** | 1.000 | 0.998 | 0.532 | 0.526 | 0.813 | 0.525 | **0.531** | 0.715 | 0.529 | 1.000 | 0.534 |
| **8HZ4** | 0.535 | 0.541 | 0.502 | 0.503 | 0.550 | 0.503 | **0.524** | 0.509 | 0.510 | 0.534 | 1.000 |

B

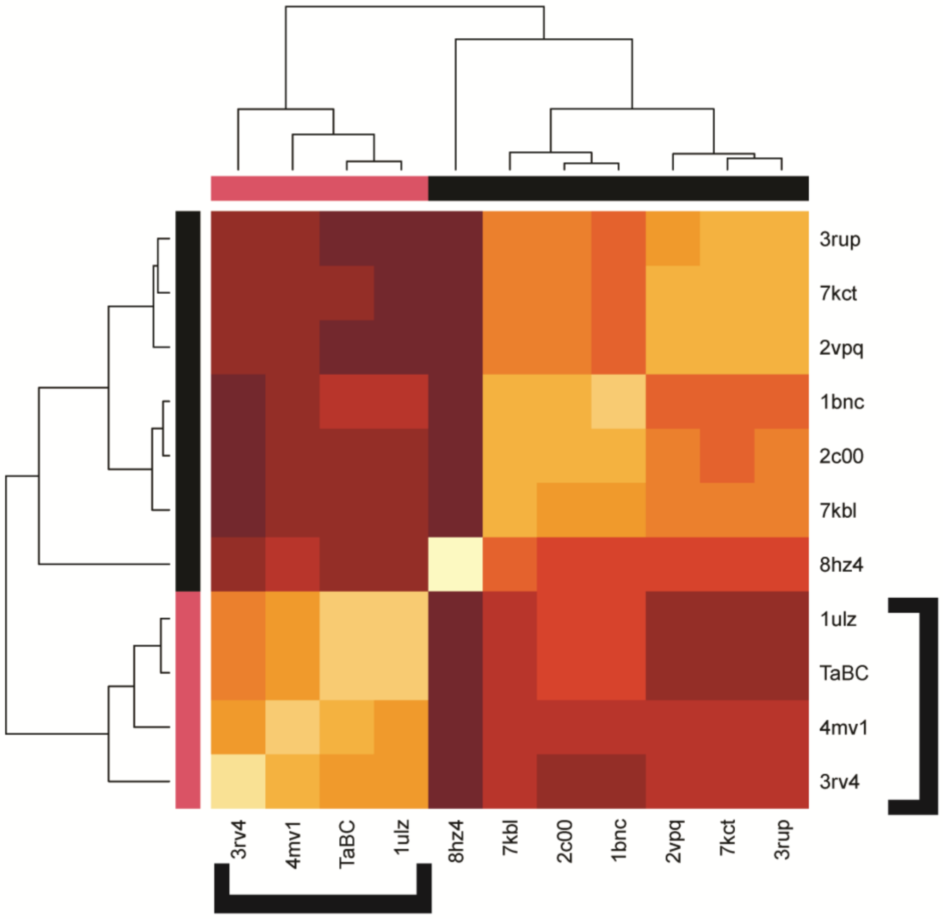


Figure S3. Comparisons of experimental structures of BC domains in terms of A) sequence identity and B) principal component analysis. The cluster with pennycress (*T. arvense*) BC is shown with brackets. The four-digit PDB accession code is listed on the axes. The identity of each protein is listed at RCSB and in Fig. 2.

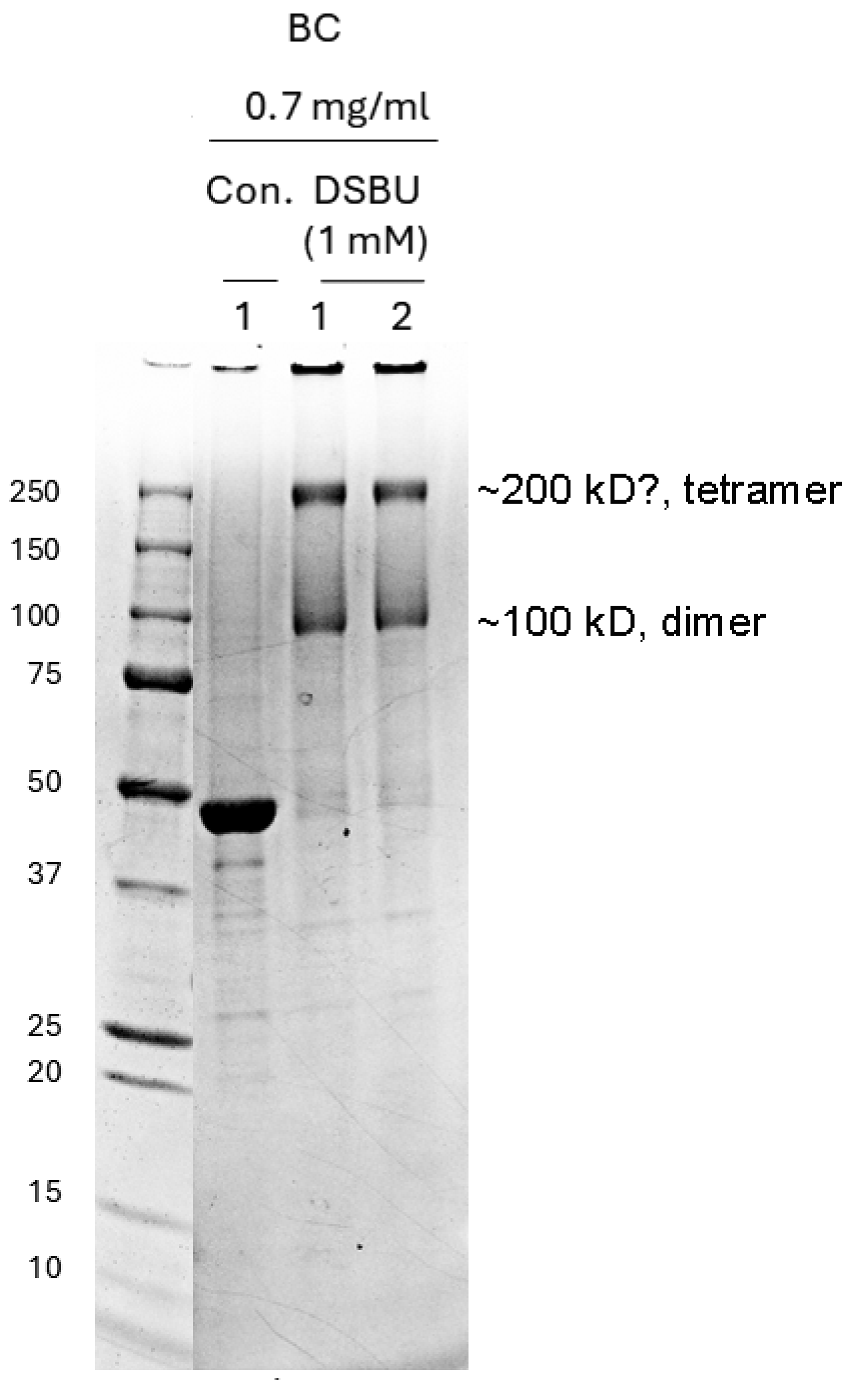


Figure S4. Cross-linking of pennycress BC with mass-spec-cleavable DSBU generated bands appearing to be dimers and tetramers on SDS-PAGE.

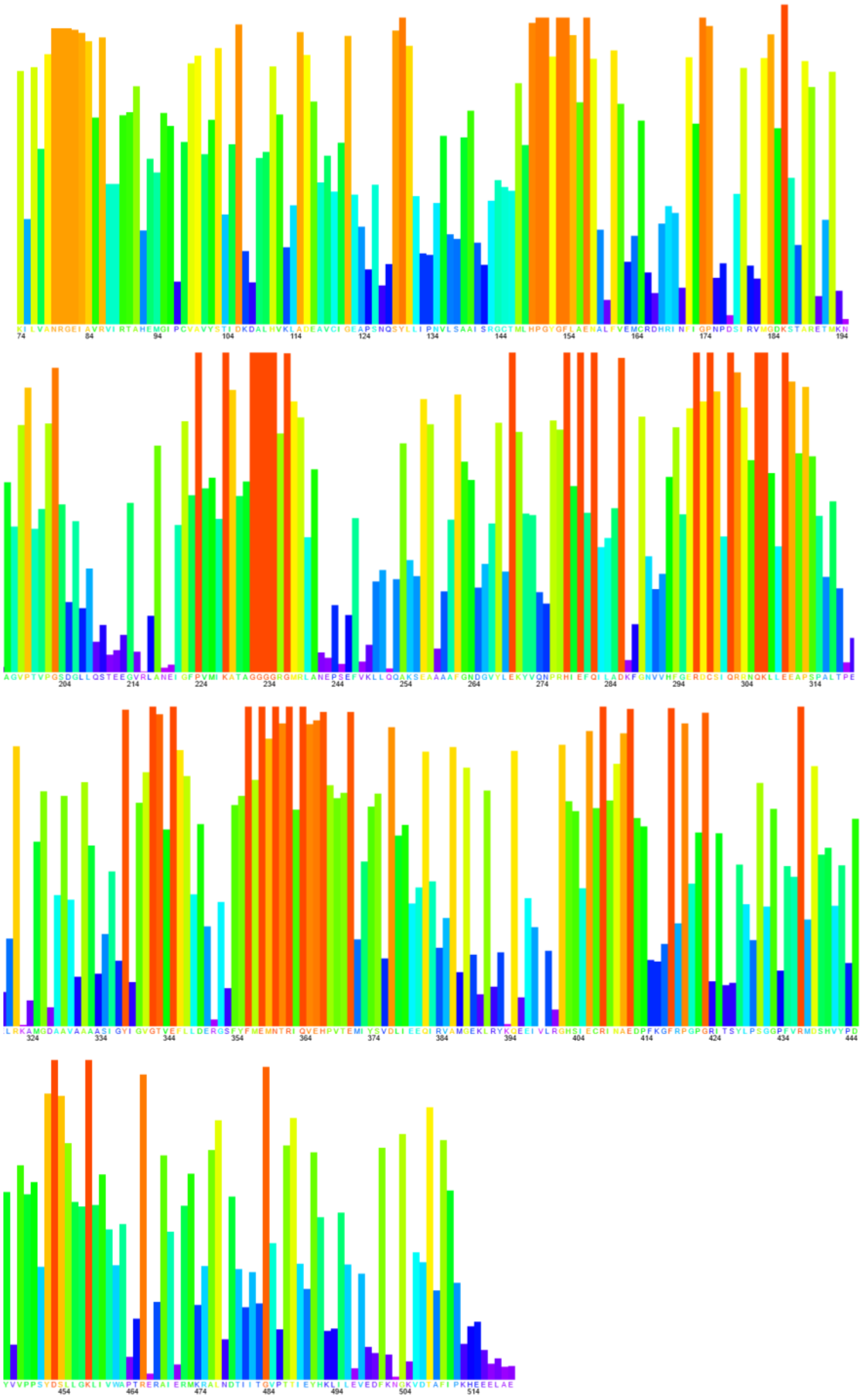

Figure S5. The residue importance from rvET, a quantification of conservation, was plotted across the sequence of mature pennycress BC. The height, on a scale of 0 to 1, marks the residue importance by rvET using 500 non-redundant sequences of BC. The color scale is analogous to that of Fig. S3 with red-orange marking the most important (conserved) and blue the least important positions in sequence.

Table S1. Cryo-EM data collection, refinement and validation statistics

Table S2. List of 500 unique homologues to pennycress BC producing significant alignments in the MSA used in the rvET analysis.

Table S3. Evolutionary importance of each sequence position of *Brassica* BC. Lower coverage and lower score implies higher functional or structural importance.

Tabel S4. Summary of DSBU crosslinks in pennycress BC, in at least two sets of measurements.

Table S5. Summary of mass spectroscopy data for acetylation assay
