## Supplemental Table 1 for "Oligomeric assemblies of plant biotin carboxylase revealed by cryo-EM and cross-linking"

**Cryo-EM data collection, refinement and validation statistics**

|  | **BC Dimer**  (EMDB-xxxx)  (PDB xxxx) | **BC Tetramer**  (EMDB-xxxx) |
| --- | --- | --- |
| **Data collection and processing** |  |  |
| Magnification | 130,000 | |
| Voltage (kV) | 300 | |
| Electron exposure (e–/Å^2^) | 40 | |
| Defocus range (μm) | -0.3 to -3.0 | |
| Pixel size (Å) | 0.97 | |
| Symmetry imposed | C2 | C2 |
| Initial particle images (no.) | 700,000 |  |
| Final particle images (no.) | 142,226 |  |
| Masked Map resolution (Å)  Unmasked map resolution (Å)  FSC threshold | 2.85  3.3  0.143 | 6.62  17  0.143 |
| Map resolution range (Å) |  |  |
| **Refinement** |  |  |
| Initial model used | AlphaFold 3 | N/A |
| Map sharpening *B* factor (Å^2^) | -91 | -263.4 |
| **Model composition**  Non-hydrogen atoms  Protein residues  Ligands | 6,962  898  0 | N/A |
| *B* factors (Å^2^) |  |  |
| R.m.s. deviations  Bond lengths (Å)  Bond angles (°) | 0/7102  0/7102  0/7102 | N/A |
| **Validation**  MolProbity score  Clashscore  Poor rotamers (%) | 4  1.6 | N/A |
| **Ramachandran plot**  Favored (%)  Allowed (%)  Disallowed (%) | 95  5  0 | N/A |
