## Supplemental Table 2 for "Oligomeric assemblies of plant biotin carboxylase revealed by cryo-EM and cross-linking"

**Table S2. Sequences producing significant alignments.**

BLASTP 2.2.24+ used the UniRef90 database, with each entry a cluster of ≥ 90% identical sequences.

```
Score    E                                                                   (Bits)  Value
lcl|M4ESC7  Biotin carboxylase,unchar n=3 Tax=Brassica RepID=M4ESC7... 903    0.0
lcl|O04983  Biotin carboxylase, chloroplastic n=7 Tax=Brassicacea...   876    0.0
lcl|M4DTW0  Uncharacterized protein n=13 Tax=Brassica RepID=M4DTW...   876    0.0
lcl|B9HBA8  Biotin carboxylase 1, chloroplastic n=10 Tax=Pentapet...   869    0.0
lcl|V4U0R4  Uncharacterized protein n=2 Tax=Pentapetalae RepID=V4...   863    0.0
lcl|O23960  Acetyl-CoA carboxylase n=4 Tax=Papilionoideae RepID=O...   863    0.0
lcl|A0A022R4R3  Uncharacterized protein n=1 Tax=Erythranthe gutta...   858    0.0
lcl|G7LIV6  Acetyl-CoA carboxylase biotin carboxylase subunit n=5...   857    0.0
lcl|B9N843  Biotin carboxylase 2, chloroplastic n=8 Tax=fabids Re...   857    0.0
lcl|E6Y9A4  Biotin carboxylase n=1 Tax=Arachis hypogaea RepID=E6Y...   857    0.0
lcl|A0A059CS83  Uncharacterized protein n=2 Tax=Eucalyptus grandi...   854    0.0
lcl|M5X110  Uncharacterized protein n=13 Tax=Rosaceae RepID=M5X11...   853    0.0
lcl|V7CS94  Uncharacterized protein n=1 Tax=Phaseolus vulgaris Re...   842    0.0
lcl|L0HSU3  Biotin carboxylase n=1 Tax=Persea americana RepID=L0H...   839    0.0
lcl|M0TTB7  Uncharacterized protein n=4 Tax=Musa acuminata subsp....   834    0.0
lcl|W1PR82  Uncharacterized protein n=2 Tax=Amborella trichopoda ...   834    0.0
lcl|A9RGQ0  Predicted protein n=2 Tax=Physcomitrella patens subsp...   815    0.0
lcl|D2CFM8  Acetyl-CoA carboxylase n=4 Tax=Arecaceae RepID=D2CFM8...   795    0.0
lcl|D8QXH7  Putative uncharacterized protein n=2 Tax=Selaginella ...   792    0.0
lcl|A9RGP9  Predicted protein n=1 Tax=Physcomitrella patens subsp...   734    0.0
lcl|I0Z7E6  Uncharacterized protein n=1 Tax=Coccomyxa subellipsoi...   678    0.0
lcl|D0V0B6  Biotin carboxylase n=1 Tax=Chromochloris zofingiensis...   662    0.0
lcl|D8UF54  Putative uncharacterized protein n=1 Tax=Volvox carte...   660    0.0
lcl|A0A087SN66  Biotin carboxylase, chloroplastic n=1 Tax=Auxenoc...   652    0.0
lcl|U7DYC7  Uncharacterized protein n=6 Tax=rosids RepID=U7DYC7_P...   641    0.0
lcl|B4VQU4  Acetyl-CoA carboxylase, biotin carboxylase n=1 Tax=Co...   635    3e-180
lcl|Q06862  Biotin carboxylase n=26 Tax=Cyanobacteria RepID=ACCC_...   628    4e-178
lcl|L8MWV8  Acetyl-CoA carboxylase, biotin carboxylase n=1 Tax=Ps...   624    9e-177
lcl|U9W9G7  Acetyl-carboxylase subunit alpha n=2 Tax=Leptolyngbya...   620    1e-175
lcl|M1WWX8  Biotin carboxylase of acetyl-CoA carboxylase n=3 Tax=...   617    8e-175
lcl|D8G4J4  Acetyl-CoA carboxylase biotin carboxylase subunit n=1...   608    4e-172
lcl|A0A077JD79  Acetyl-CoA carboxylase biotin carboxylase subunit...   608    4e-172
lcl|G4FNX9  Acetyl-CoA carboxylase, biotin carboxylase n=1 Tax=Sy...   605    4e-171
lcl|L8M6P6  Acetyl-CoA carboxylase, biotin carboxylase subunit n=...   602    4e-170
lcl|B4WQC2  Acetyl-CoA carboxylase, biotin carboxylase n=1 Tax=Sy...   601    6e-170
lcl|U5DND4  Acetyl-CoA carboxylase, biotin carboxylase subunit n=...   600    8e-170
lcl|A0A059LN48  L chain of carbamoyl-phosphate synthase n=1 Tax=H...   598    5e-169
lcl|B5ILR5  Acetyl-CoA carboxylase, biotin carboxylase n=3 Tax=Ch...   593    1e-167
lcl|B8LR39  Putative uncharacterized protein n=1 Tax=Picea sitche...   591    7e-167
lcl|A8JGF4  Biotin carboxylase, acetyl-CoA carboxylase component ...   585    3e-165
lcl|L8LM23  Acetyl-CoA carboxylase, biotin carboxylase subunit n=...   579    2e-163
lcl|B1X573  Acetyl-CoA carboxylase n=1 Tax=Paulinella chromatopho...   575    3e-162
lcl|R7QSG8  Stackhouse genomic scaffold, scaffold_62 n=1 Tax=Chon...   560    1e-157
lcl|R7RU12  Biotin carboxylase of acetyl-CoA carboxylase n=2 Tax=...   555    6e-156
lcl|M1Z8S4  Acetyl-CoA carboxylase subunit (Biotin carboxylase su...   553    1e-155
lcl|L5MXM1  Biotin carboxylase n=6 Tax=Brevibacillus RepID=L5MXM1...   545    3e-153
lcl|K2F0J0  Uncharacterized protein n=1 Tax=uncultured bacterium ...   539    3e-151
lcl|A0A096DN89  Acetyl-CoA carboxylase n=1 Tax=Caloranaerobacter ...   538    4e-151
lcl|K8E080  Biotin carboxylase 1 n=1 Tax=Desulfotomaculum hydroth...   538    6e-151
lcl|A0A078KQG6  Biotin carboxylase 1 n=1 Tax=[Clostridium] cellul...   531    8e-149
lcl|U1X0I3  Acetyl-CoA carboxylase, biotin carboxylase subunit n=...   527    9e-148
lcl|M3EEM0  Biotin carboxylase of acetyl-CoA carboxylase n=4 Tax=...   526    2e-147
lcl|B4D215  Acetyl-CoA carboxylase, biotin carboxylase n=1 Tax=Ch...   525    4e-147
lcl|A0A069DHM6  Biotin carboxylase n=1 Tax=Paenibacillus sp. TCA2...   524    9e-147
lcl|A0A087LFX5  Acetyl-CoA carboxylase n=22 Tax=Geobacillus RepID...   523    2e-146
lcl|A0A094JPF0  Acetyl-CoA carboxylase n=1 Tax=Thermoactinomyces ...   523    2e-146
lcl|T0C7L7  Acetyl-CoA carboxylase biotin carboxylase subunit n=1...   521    8e-146
lcl|A0A074LRS0  Acetyl-CoA carboxylase n=1 Tax=Tumebacillus flage...   520    1e-145
lcl|A0A073JYA5  Acetyl-CoA carboxylase n=1 Tax=Bacillus manlipone...   519    3e-145
lcl|H3SFE9  Acetyl-CoA carboxylase, biotin carboxylase n=3 Tax=Pa...   518    4e-145
lcl|G9QPK3  Biotin carboxylase 1 n=1 Tax=Bacillus smithii 7_3_47F...   518    7e-145
lcl|D4W2R2  Acetyl-CoA carboxylase, biotin carboxylase subunit n=...   516    2e-144
lcl|S2YPG1  Biotin carboxylase 1 n=4 Tax=Planococcaceae RepID=S2Y...   515    5e-144
lcl|R9C238  Acetyl-CoA carboxylase biotin carboxylase subunit n=2...   514    8e-144
lcl|I9M443  Acetyl-CoA carboxylase, biotin carboxylase n=3 Tax=Pe...   514    9e-144
lcl|A0A081SE02  Acetyl-CoA carboxylase n=1 Tax=Chlorobium sp. GBC...   514    1e-143
lcl|I8UKH6  Acetyl-CoA carboxylase biotin carboxylase subunit n=1...   514    1e-143
lcl|A0A061PA86  Biotin carboxylase of acetyl-CoA carboxylase n=2 ...   514    1e-143
lcl|F5LLQ6  Acetyl-CoA carboxylase, biotin carboxylase subunit n=...   514    1e-143
lcl|S9UU84  Acetyl-CoA carboxylase, biotin carboxylase n=3 Tax=Pa...   513    2e-143
lcl|F5SB10  Acetyl-CoA carboxylase, biotin carboxylase n=1 Tax=De...   513    2e-143
lcl|P49787  Biotin carboxylase 1 n=69 Tax=Bacillus RepID=ACCC1_BACSU   512    5e-143
lcl|R9NDB8  Acetyl-CoA carboxylase, biotin carboxylase subunit n=...   511    7e-143
lcl|N4WAW6  Acetyl-CoA carboxylase biotin carboxylase subunit n=1...   511    8e-143
lcl|A0A084GUF2  Acetyl-CoA carboxylase n=2 Tax=Bacillus RepID=A0A...   511    8e-143
lcl|J0LET8  Acetyl-CoA carboxylase, biotin carboxylase n=3 Tax=Po...   511    8e-143
lcl|I8TN07  Acetyl-CoA carboxylase, biotin carboxylase n=1 Tax=Pe...   511    1e-142
lcl|A0A024QAY3  Biotin carboxylase n=3 Tax=Bacillaceae RepID=A0A0...   511    1e-142
lcl|U2E8F2  Acetyl-CoA carboxylase biotin carboxylase subunit pro...   510    2e-142
lcl|A0A074L558  Acetyl-CoA carboxylase n=1 Tax=Anditalea andensis...   509    3e-142
lcl|M7NWA7  Biotin carboxylase n=1 Tax=Cesiribacter andamanensis ...   508    5e-142
lcl|M1V7A2  Biotin carboxylase, chloroplast n=1 Tax=Cyanidioschyz...   508    7e-142
lcl|A0A015P683  Acetyl-CoA carboxylase n=1 Tax=Paenibacillus darw...   508    8e-142
lcl|A0A081P467  Acetyl-CoA carboxylase n=2 Tax=Paenibacillus RepI...   508    9e-142
lcl|A0A037YZI5  Biotin carboxylase n=1 Tax=Clostridium tetanomorp...   507    1e-141
lcl|H8I9A4  Putative pyruvate carboxylase subunit A n=1 Tax=Metha...   506    2e-141
lcl|Q1K2Y2  Acetyl-CoA carboxylase, biotin carboxylase n=1 Tax=De...   506    3e-141
lcl|W2UNB6  Acetyl-CoA carboxylase, biotin carboxylase subunit n=...   504    7e-141
lcl|M2Q2W3  Acetyl-CoA carboxylase, biotin carboxylase subunit n=...   504    7e-141
lcl|B1QYW6  Acetyl-CoA carboxylase, biotin carboxylase n=6 Tax=Cl...   504    9e-141
lcl|V6SV80  Acetyl-CoA carboxylase biotin carboxylase subunit n=2...   504    1e-140
lcl|W9BYT4  Acetyl-CoA carboxylase biotin carboxylase subunit n=2...   504    1e-140
lcl|W7YWA8  Biotin carboxylase of acetyl-CoA carboxylase n=1 Tax=...   504    1e-140
lcl|K2Q7I8  Acetyl-CoA carboxylase, biotin carboxylase n=3 Tax=Fl...   503    2e-140
lcl|E7GII1  Uncharacterized protein n=4 Tax=Clostridiales RepID=E...   503    2e-140
lcl|A0A090ZGU1  Acetyl-CoA carboxylase, biotin carboxylase subuni...   503    3e-140
lcl|E0ICJ1  Acetyl-CoA carboxylase, biotin carboxylase n=1 Tax=Pa...   503    3e-140
lcl|A1ZN48  Acetyl-CoA carboxylase, biotin carboxylase n=1 Tax=Mi...   501    1e-139
lcl|W8YM74  Biotin carboxylase 1 n=3 Tax=Paenibacillus RepID=W8YM...   501    1e-139
lcl|W7CHS6  Acetyl-CoA carboxylase biotin carboxylase subunit n=3...   500    2e-139
lcl|D1YUY4  Putative pyruvate carboxylase subunit A n=1 Tax=Metha...   500    2e-139
lcl|M7NHG0  Biotin carboxylase n=1 Tax=Bhargavaea cecembensis DSE...   499    2e-139
lcl|D7V015  Acetyl-CoA carboxylase, biotin carboxylase subunit n=...   499    3e-139
lcl|H7F2S2  Acetyl-CoA carboxylase biotin carboxylase subunit n=4...   499    3e-139
lcl|V9HH99  Acetyl-CoA carboxylase, biotin carboxylase subunit n=...   499    3e-139
lcl|R9MRR7  Acetyl-CoA carboxylase, biotin carboxylase subunit n=...   498    5e-139
lcl|U4P3P9  Biotin carboxylase n=5 Tax=Clostridium botulinum RepI...   498    5e-139
lcl|Q5NVT4  Pyruvate carboxylase subunit a n=2 Tax=Archaea RepID=...   498    6e-139
lcl|I0R4Y0  Acetyl-CoA carboxylase, biotin carboxylase subunit n=...   498    1e-138
lcl|X0RIU1  Biotin carboxylase n=1 Tax=Bacillus sp. TS-2 RepID=X0...   497    1e-138
lcl|O27939  Pyruvate carboxylase subunit A n=3 Tax=Methanothermob...   497    1e-138
lcl|S2YUY5  Acetyl-CoA carboxylase, biotin carboxylase subunit n=...   496    2e-138
lcl|M5U9Y7  Acetyl-CoA carboxylase biotin carboxylase subunit n=2...   496    3e-138
lcl|A0A069D968  Biotin carboxylase of acetyl-CoA carboxylase n=1 ...   496    4e-138
lcl|U2NPN3  Biotin carboxylase n=1 Tax=Clostridium intestinale UR...   495    4e-138
lcl|N2AUT7  Acetyl-CoA carboxylase, biotin carboxylase subunit n=...   495    6e-138
lcl|A0A023BZD7  Acetyl-CoA carboxylase n=4 Tax=Aquimarina RepID=A...   494    6e-138
lcl|B1CA69  Acetyl-CoA carboxylase, biotin carboxylase subunit n=...   494    8e-138
lcl|C0BIJ7  Acetyl-CoA carboxylase, biotin carboxylase n=1 Tax=Fl...   494    9e-138
lcl|K2GPI5  Acetyl-CoA carboxylase biotin carboxylase subunit n=1...   494    1e-137
lcl|U1FRU5  Acetyl-CoA carboxylase biotin carboxylase subunit n=2...   494    1e-137
lcl|W4VDI5  Biotin carboxylase of acetyl-CoA carboxylase n=1 Tax=...   493    2e-137
lcl|S6EQK6  Putative Acetyl-CoA carboxylase, biotin carboxylase s...   493    2e-137
lcl|K4MLR7  Acetyl-CoA carboxylase biotin carboxylase n=1 Tax=Met...   493    2e-137
lcl|T0IDC6  Biotin carboxylase 1 n=1 Tax=Sporomusa ovata DSM 2662...   493    2e-137
lcl|C6PQ77  Acetyl-CoA carboxylase, biotin carboxylase n=3 Tax=Cl...   493    3e-137
lcl|S9R3B6  Acetyl-CoA carboxylase, biotin carboxylase subunit n=...   492    3e-137
lcl|A0A095CJJ5  Acetyl-CoA carboxylase n=1 Tax=Rhodovulum sp. NI2...   492    4e-137
lcl|B1QZJ5  Acetyl-CoA carboxylase, biotin carboxylase n=6 Tax=Cl...   491    8e-137
lcl|F4MMI1  Acetyl-CoA carboxylase biotin carboxylase subunit n=1...   491    8e-137
lcl|F0YTH2  Acetyl-CoA carboxylase, biotin carboxylase subunit n=...   491    1e-136
lcl|A0A078MFB4  Biotin carboxylase n=2 Tax=Lysinibacillus RepID=A...   490    1e-136
lcl|A0A058ZJL4  Acetyl-CoA carboxylase biotin carboxylase subunit...   490    1e-136
lcl|W4ENP6  Acetyl-CoA carboxylase biotin carboxylase subunit n=1...   490    2e-136
lcl|F0IEV0  Acetyl-CoA carboxylase, biotin carboxylase n=5 Tax=Ca...   490    2e-136
lcl|N1ZSB0  Acetyl-CoA carboxylase, biotin carboxylase subunit n=...   490    2e-136
lcl|A0A086AUE7  Acetyl-CoA carboxylase n=1 Tax=Flavobacterium hyd...   489    2e-136
lcl|Y5P1R4  Acetyl-CoA carboxylase, biotin carboxylase subunit n=...   489    3e-136
lcl|D7DR99  Acetyl-CoA carboxylase, biotin carboxylase n=1 Tax=Me...   488    6e-136
lcl|H2BTR3  Acetyl-CoA carboxylase, biotin carboxylase n=2 Tax=Gi...   488    6e-136
lcl|A0A085L124  Acetyl-CoA carboxylase subunit alpha n=1 Tax=Schl...   488    8e-136
lcl|A0A063Z2K6  Acetyl-CoA carboxylase n=12 Tax=Bacillus RepID=A0...   488    9e-136
lcl|U2SQ79  Biotin carboxylase of acetyl-CoA carboxylase protein ...   487    2e-135
lcl|U2UY92  Uncharacterized protein n=2 Tax=Leptotrichia RepID=U2...   486    2e-135
lcl|G5INN8  Acetyl-CoA carboxylase, biotin carboxylase n=1 Tax=[C...   486    2e-135
lcl|G5GKY5  Acetyl-CoA carboxylase n=1 Tax=Johnsonella ignava ATC...   486    2e-135
lcl|C3WEY9  Acetyl-CoA carboxylase, biotin carboxylase subunit n=...   486    2e-135
lcl|U4KQJ2  Acetyl-CoA carboxylase, biotin carboxylase n=1 Tax=Ac...   486    3e-135
lcl|A0A031IAA6  Acetyl-CoA carboxylase, biotin carboxylase n=6 Ta...   486    3e-135
lcl|U2XXC5  Fumarate reductase flavoprotein subunit n=2 Tax=uncla...   485    4e-135
lcl|F4BTD8  Acetyl-CoA carboxylase n=1 Tax=Methanosaeta concilii ...   485    7e-135
lcl|M2TA93  Biotin carboxylase of acetyl-CoA carboxylase n=1 Tax=...   484    7e-135
lcl|D7E897  Acetyl-CoA carboxylase, biotin carboxylase n=1 Tax=Me...   484    1e-134
lcl|A0A060QGJ2  Biotin carboxylase of acetyl-CoA carboxylase n=1 ...   484    1e-134
lcl|S9SH73  Biotin carboxylase of acetyl-CoA carboxylase n=1 Tax=...   483    2e-134
lcl|A0A072X6Z0  Acetyl-CoA carboxylase n=2 Tax=Clostridium botuli...   483    2e-134
lcl|W9AGW3  Biotin carboxylase n=1 Tax=Oceanobacillus picturae Re...   483    2e-134
lcl|W6K8T0  Acetyl-CoA carboxylase, biotin carboxylase subunit n=...   483    3e-134
lcl|K2HS24  Acetyl-CoA carboxylase n=1 Tax=Oceaniovalibus guishan...   482    4e-134
lcl|K2IFX7  Acetyl-CoA carboxylase biotin carboxylase subunit n=1...   482    4e-134
lcl|A0A072NKD8  Acetyl-CoA carboxylase, biotin carboxylase subuni...   482    5e-134
lcl|A0A077VQE9  Biotin carboxylase n=4081 Tax=Staphylococcus RepI...   482    5e-134
lcl|F0TB64  Acetyl-CoA carboxylase, biotin carboxylase n=1 Tax=Me...   482    5e-134
lcl|Q12TE9  Pyruvate carboxylase subunit A n=1 Tax=Methanococcoid...   481    6e-134
lcl|J9HED6  Acetyl-CoA carboxylase, biotin carboxylase n=1 Tax=Al...   481    8e-134
lcl|A0A037ZHZ4  Acetyl-CoA carboxylase n=1 Tax=Actibacterium muco...   481    1e-133
lcl|A3W8A6  Acetyl-CoA carboxylase n=2 Tax=Roseovarius RepID=A3W8...   481    1e-133
lcl|A3TT03  Acetyl-CoA carboxylase n=1 Tax=Oceanicola batsensis (...   479    2e-133
lcl|A5ZAG3  Acetyl-CoA carboxylase, biotin carboxylase subunit n=...   479    2e-133
lcl|K8Z3B6  Acetyl-CoA carboxylase, biotin carboxylase subunit n=...   479    2e-133
lcl|D5EAE6  Pyruvate carboxylase subunit A n=1 Tax=Methanohalophi...   479    2e-133
lcl|A0B7A9  Pyruvate carboxylase subunit A n=1 Tax=Methanosaeta t...   479    3e-133
lcl|W1UUU3  Uncharacterized protein n=1 Tax=Intestinibacter bartl...   479    4e-133
lcl|S0JGC2  Acetyl-CoA carboxylase, biotin carboxylase subunit n=...   479    4e-133
lcl|V7I5I1  Acetyl-CoA carboxylase biotin carboxylase subunit n=1...   479    5e-133
lcl|K2JEN3  Acetyl-CoA carboxylase biotin carboxylase subunit n=1...   478    5e-133
lcl|G9YL09  Acetyl-CoA carboxylase, biotin carboxylase subunit n=...   478    7e-133
lcl|A0A085KB10  Acetyl-CoA carboxylase n=18 Tax=cellular organism...   478    8e-133
lcl|F7JIZ0  Acetyl-CoA carboxylase, biotin carboxylase n=1 Tax=La...   478    9e-133
lcl|D0WFZ6  Acetyl-CoA carboxylase, biotin carboxylase subunit n=...   478    9e-133
lcl|A0A090I4J0  Pyruvate carboxylase subunit A n=4 Tax=Methanobac...   478    9e-133
lcl|Q2NFV1  PycA n=1 Tax=Methanosphaera stadtmanae (strain ATCC 4...   478    1e-132
lcl|H1PV87  Acetyl-CoA carboxylase, biotin carboxylase subunit n=...   477    1e-132
lcl|F5RCS6  Acetyl-CoA carboxylase subunit A n=4 Tax=Rhodocyclale...   477    1e-132
lcl|T0L8E1  Acetyl-CoA carboxylase, biotin carboxylase n=1 Tax=ca...   477    1e-132
lcl|Q8PVX7  Pyruvate carboxylase, subunit A n=6 Tax=Methanosarcin...   477    1e-132
lcl|A0A088R6E2  Acetyl-CoA carboxylase n=300 Tax=Proteobacteria R...   476    2e-132
lcl|I4EGQ5  Acetyl-CoA carboxylase, biotin carboxylase subunit n=...   476    2e-132
lcl|Q0FI32  Acetyl-CoA carboxylase n=3 Tax=Rhodobacteraceae RepID...   476    2e-132
lcl|K1Y9U0  Uncharacterized protein n=1 Tax=uncultured bacterium ...   476    3e-132
lcl|F7RXF9  Acetyl-CoA carboxylase, biotin carboxylase subunit n=...   476    3e-132
lcl|A3V3V2  Acetyl-CoA carboxylase n=2 Tax=Loktanella vestfoldens...   475    4e-132
lcl|J1FBH7  Acetyl-CoA carboxylase, biotin carboxylase subunit n=...   475    6e-132
lcl|R9SNA6  Pyruvate carboxylase subunit A PycA n=1 Tax=Methanobr...   474    7e-132
lcl|A6URX6  Acetyl-CoA carboxylase, biotin carboxylase n=1 Tax=Me...   474    1e-131
lcl|G5ZXZ8  Acetyl-CoA carboxylase, biotin carboxylase n=1 Tax=SA...   474    1e-131
lcl|I5C1C6  Acetyl-CoA carboxylase biotin carboxylase subunit n=4...   474    1e-131
lcl|L0KUC7  Acetyl-CoA carboxylase, biotin carboxylase subunit n=...   474    1e-131
lcl|A5UL92  Pyruvate carboxylase (Acetyl-CoA/biotin carboxylase),...   473    2e-131
lcl|D6TRA5  Acetyl-CoA carboxylase, biotin carboxylase n=1 Tax=Kt...   473    2e-131
lcl|E0WT96  Biotin carboxylase n=1 Tax=Candidatus Regiella insect...   473    2e-131
lcl|L1NUR8  Acetyl-CoA carboxylase, biotin carboxylase subunit n=...   473    3e-131
lcl|D3E2D7  Pyruvate carboxylase subunit A PycA n=1 Tax=Methanobr...   472    4e-131
lcl|D5VR94  Acetyl-CoA carboxylase, biotin carboxylase n=1 Tax=Me...   472    4e-131
lcl|Q0EWA0  Acetyl-CoA carboxylase n=2 Tax=Mariprofundus RepID=Q0...   472    4e-131
lcl|A0A059IUJ8  Acetyl-CoA carboxylase biotin carboxylase subunit...   472    5e-131
lcl|H5SM68  Acetyl-CoA carboxylase, biotin carboxylase subunit n=...   472    5e-131
lcl|W9DZ19  Pyruvate carboxylase subunit A n=1 Tax=Methanolobus t...   472    6e-131
lcl|F6D7M1  Acetyl-CoA carboxylase, biotin carboxylase n=1 Tax=Me...   471    6e-131
lcl|E0MKE6  Acetyl-CoA carboxylase, biotin carboxylase subunit n=...   471    7e-131
lcl|S3XM00  Acetyl-CoA carboxylase, biotin carboxylase subunit n=...   471    7e-131
lcl|G5HBM0  Acetyl-CoA carboxylase, biotin carboxylase n=1 Tax=Al...   471    8e-131
lcl|A0A085FM63  Biotin carboxylase of acetyl-CoA carboxylase n=2 ...   471    9e-131
lcl|G0H2N8  Pyruvate carboxylase subunit A n=7 Tax=Methanococcus ...   471    1e-130
lcl|A3XRD6  Acetyl-CoA carboxylase, biotin carboxylase n=2 Tax=Le...   470    1e-130
lcl|I0JXW1  Pyruvate carboxylase subunit A n=1 Tax=Methylacidiphi...   470    1e-130
lcl|A4BP25  Acetyl-CoA carboxylase n=1 Tax=Nitrococcus mobilis Nb...   470    1e-130
lcl|J0QZQ7  Acetyl-CoA carboxylase, biotin carboxylase subunit n=...   470    1e-130
lcl|V8A4D2  Acetyl-CoA carboxylase biotin carboxylase subunit n=7...   470    2e-130
lcl|K0Z8H3  Acetyl-CoA carboxylase, biotin carboxylase subunit n=...   470    2e-130
lcl|X5MP46  Biotin carboxylase of acetyl-CoA carboxylase n=1 Tax=...   470    2e-130
lcl|H5SD30  Acetyl-CoA carboxylase n=1 Tax=uncultured gamma prote...   470    2e-130
lcl|Q58626  Pyruvate carboxylase subunit A n=5 Tax=Methanocaldoco...   470    2e-130
lcl|U2DRE7  Acetyl-CoA carboxylase, biotin carboxylase subunit n=...   469    2e-130
lcl|N2JA18  Biotin carboxylase n=5 Tax=Pseudomonadaceae RepID=N2J...   469    4e-130
lcl|S7J5J6  Acetyl-CoA carboxylase, biotin carboxylase subunit n=...   468    5e-130
lcl|W7QEE2  Biotin carboxylase n=3 Tax=Alteromonadaceae RepID=W7Q...   468    5e-130
lcl|A0A062V4F0  Acetyl-CoA carboxylase, biotin carboxylase subuni...   468    6e-130
lcl|J2IHY9  Acetyl-CoA carboxylase biotin carboxylase subunit n=8...   468    6e-130
lcl|S9PFM3  Biotin carboxylase of acetyl-CoA carboxylase n=2 Tax=...   468    8e-130
lcl|R9LI17  Acetyl-CoA carboxylase, biotin carboxylase subunit n=...   468    9e-130
lcl|A0A074JX28  Acetyl-CoA carboxylase biotin carboxylase subunit...   468    1e-129
lcl|K2DX19  Uncharacterized protein n=1 Tax=uncultured bacterium ...   467    1e-129
lcl|R9KIT9  Acetyl-CoA carboxylase, biotin carboxylase subunit n=...   467    1e-129
lcl|A0A060NPT1  Biotin carboxylase n=3 Tax=unclassified Comamonad...   466    2e-129
lcl|G7WLD2  Pyruvate carboxylase subunit A n=2 Tax=Methanosaeta h...   466    2e-129
lcl|P37798  Biotin carboxylase n=222 Tax=Pseudomonas RepID=ACCC_P...   466    3e-129
lcl|A0A086YAZ5  Acetyl-CoA carboxylase n=2 Tax=Haematobacter RepI...   466    3e-129
lcl|A0A094IQ85  Acetyl-CoA carboxylase n=1 Tax=Idiomarina sp. MCC...   466    4e-129
lcl|D4TU10  Acetyl-CoA carboxylase, biotin carboxylase n=1 Tax=Ra...   465    4e-129
lcl|A0A085Y240  Biotin carboxylase of acetyl-CoA carboxylase n=1 ...   465    5e-129
lcl|J0S071  Acetyl-CoA carboxylase, biotin carboxylase n=1 Tax=Me...   465    6e-129
lcl|A0A066TCN5  Biotin carboxylase of acetyl-CoA carboxylase n=4 ...   465    6e-129
lcl|T4V4D1  Acetyl-CoA carboxylase, biotin carboxylase subunit n=...   464    8e-129
lcl|J1H614  Acetyl-CoA carboxylase, biotin carboxylase subunit n=...   464    8e-129
lcl|D3DJ42  2-oxoglutarate carboxylase small subunit n=4 Tax=Aqui...   464    8e-129
lcl|E4L2B3  Acetyl-CoA carboxylase, biotin carboxylase subunit n=...   464    1e-128
lcl|N6V1P2  Pyruvate carboxylase subunit A n=1 Tax=Methanocaldoco...   464    1e-128
lcl|T1D1D1  Biotin carboxylase of acetyl-CoA carboxylase n=1 Tax=...   464    1e-128
lcl|F8ALC9  Acetyl-CoA carboxylase, biotin carboxylase n=1 Tax=Me...   463    2e-128
lcl|I8I2I8  Acetyl-CoA carboxylase, biotin carboxylase n=1 Tax=Hy...   463    2e-128
lcl|Q1Q0S1  Strongly similar to biotin carboxylase (A subunit of ...   462    3e-128
lcl|E6MHW6  Acetyl-CoA carboxylase, biotin carboxylase subunit n=...   462    3e-128
lcl|A6GHC5  Acetyl-CoA carboxylase, biotin carboxylase n=1 Tax=Pl...   462    3e-128
lcl|E3CJN8  Acetyl-CoA carboxylase, biotin carboxylase subunit n=...   462    3e-128
lcl|A6UVR2  Acetyl-CoA carboxylase, biotin carboxylase n=1 Tax=Me...   462    4e-128
lcl|E7G8T8  Acetyl-CoA carboxylase n=2 Tax=Erysipelotrichaceae Re...   462    4e-128
lcl|A0A073D2G4  Acetyl-CoA carboxylase n=1 Tax=Clostridium botuli...   462    5e-128
lcl|U2UUB2  Acetyl-CoA carboxylase, biotin carboxylase subunit n=...   462    5e-128
lcl|G9WMZ4  Acetyl-CoA carboxylase n=4 Tax=Oribacterium RepID=G9W...   462    5e-128
lcl|I4YZS2  Acetyl-CoA carboxylase, biotin carboxylase subunit n=...   462    6e-128
lcl|E2CRT1  Acetyl-CoA carboxylase, biotin carboxylase subunit n=...   462    6e-128
lcl|A0A059KMA5  Acetyl-CoA carboxylase, biotin carboxylase n=1 Ta...   461    7e-128
lcl|F3L057  Biotin carboxylase of acetyl-CoA carboxylase n=1 Tax=...   461    7e-128
lcl|V6SI43  Biotin carboxylase n=4 Tax=Flavobacterium RepID=V6SI4...   461    8e-128
lcl|G4E1T3  Acetyl-CoA carboxylase, biotin carboxylase n=1 Tax=Th...   461    1e-127
lcl|W2BXF3  Acetyl-CoA carboxylase, biotin carboxylase subunit n=...   461    1e-127
lcl|H3KCV6  Acetyl-CoA carboxylase, biotin carboxylase subunit n=...   461    1e-127
lcl|S1NVJ0  Acetyl-CoA carboxylase, biotin carboxylase subunit n=...   460    2e-127
lcl|P43873  Biotin carboxylase n=44 Tax=Pasteurellaceae RepID=ACC...   460    2e-127
lcl|G9ZDN0  Acetyl-CoA carboxylase, biotin carboxylase subunit n=...   460    2e-127
lcl|F0EYW5  Acetyl-CoA carboxylase, biotin carboxylase subunit n=...   460    2e-127
lcl|A0A084T135  Pyruvate carboxylase subunit A n=2 Tax=Cystobacte...   459    2e-127
lcl|K1ZFE5  Uncharacterized protein n=2 Tax=uncultured bacterium ...   459    3e-127
lcl|W9H606  Acetyl-CoA carboxylase biotin carboxylase subunit n=1...   459    3e-127
lcl|P24182  Biotin carboxylase n=3788 Tax=cellular organisms RepI...   459    3e-127
lcl|A0A085FFT5  Acetyl-CoA carboxylase, biotin carboxylase subuni...   459    4e-127
lcl|J2KHP9  Acetyl/propionyl-CoA carboxylase, alpha subunit n=1 T...   459    4e-127
lcl|Q2FNH4  Pyruvate carboxylase subunit A n=1 Tax=Methanospirill...   459    5e-127
lcl|V4TG18  Biotin carboxylase of acetyl-CoA carboxylase n=1 Tax=...   458    5e-127
lcl|S0KPM8  Acetyl-CoA carboxylase, biotin carboxylase subunit n=...   458    5e-127
lcl|A0A095VNZ3  Biotin carboxylase of acetyl-CoA carboxylase n=1 ...   458    7e-127
lcl|V7IER2  Biotin carboxylase n=2 Tax=Eikenella corrodens RepID=...   458    7e-127
lcl|D4YGY1  Acetyl-CoA carboxylase, biotin carboxylase subunit n=...   458    8e-127
lcl|A0A081G436  Biotin carboxylase of acetyl-CoA carboxylase n=2 ...   458    8e-127
lcl|G4CSY8  Acetyl-CoA carboxylase, biotin carboxylase n=1 Tax=Ne...   457    9e-127
lcl|F5TD40  Acetyl-CoA carboxylase, biotin carboxylase subunit n=...   457    9e-127
lcl|W7UY17  Acetyl-CoA carboxylase biotin carboxylase subunit n=6...   457    9e-127
lcl|H0A7F1  Acetyl-CoA carboxylase, biotin carboxylase subunit n=...   457    1e-126
lcl|A0A016XMG1  Acetyl-CoA carboxylase n=2 Tax=Hylemonella gracil...   457    1e-126
lcl|C3X596  Biotin carboxylase n=2 Tax=Oxalobacter formigenes Rep...   457    1e-126
lcl|F7XQF3  Acetyl-CoA carboxylase, biotin carboxylase n=1 Tax=Me...   457    2e-126
lcl|G4CKI7  Acetyl-CoA carboxylase, biotin carboxylase n=1 Tax=Ne...   457    2e-126
lcl|E7FZE8  Biotin carboxylase n=2 Tax=Helicobacter suis RepID=E7...   456    2e-126
lcl|N6W6F8  Acetyl-CoA carboxylase, biotin carboxylase subunit n=...   456    3e-126
lcl|K5YUR1  Acetyl-CoA carboxylase biotin carboxylase subunit n=2...   456    4e-126
lcl|H1G3A8  Acetyl-CoA carboxylase, biotin carboxylase n=1 Tax=Ec...   455    5e-126
lcl|N6YRK7  Acetyl-CoA carboxylase biotin carboxylase subunit n=4...   455    5e-126
lcl|H1H1F6  Acetyl-CoA carboxylase, biotin carboxylase subunit n=...   455    7e-126
lcl|C6L8T7  Acetyl-CoA carboxylase, biotin carboxylase subunit n=...   454    7e-126
lcl|A0P4V7  Acetyl-CoA carboxylase n=1 Tax=Methylophilales bacter...   454    9e-126
lcl|A0A078LBE1  Biotin carboxylase n=1 Tax=Chlamydia sp. 'Rubis' ...   454    9e-126
lcl|D1KCB1  Biotin carboxylase n=1 Tax=uncultured SUP05 cluster b...   454    1e-125
lcl|W9V645  Biotin carboxylase n=2 Tax=Nitrincola RepID=W9V645_9GAMM   454    1e-125
lcl|O52058  Biotin carboxylase n=6 Tax=Chromatiaceae RepID=ACCC_A...   454    1e-125
lcl|A0A062XUE6  Acetyl-CoA carboxylase n=2 Tax=root RepID=A0A062X...   454    1e-125
lcl|H2BV31  Carbamoyl-phosphate synthase L chain ATP-binding n=1 ...   453    2e-125
lcl|G1Y1K3  Biotin carboxylase n=1 Tax=Nitrospirillum amazonense ...   453    2e-125
lcl|G4E777  Acetyl-CoA carboxylase, biotin carboxylase n=1 Tax=Th...   453    2e-125
lcl|N6YYA5  Methylcrotonoyl-CoA carboxylase n=1 Tax=Thauera linal...   452    3e-125
lcl|A0A084TJJ6  Biotin carboxylase n=1 Tax=Mangrovimonas yunxiaon...   452    4e-125
lcl|G2FJZ3  Biotin carboxylase n=2 Tax=sulfur-oxidizing symbionts...   452    4e-125
lcl|F6BF33  Acetyl-CoA carboxylase, biotin carboxylase n=2 Tax=Me...   452    5e-125
lcl|H1G5K0  Pyruvate carboxylase subunit A n=1 Tax=Ectothiorhodos...   452    6e-125
lcl|A0A081NKE6  Acetyl-CoA carboxylase n=1 Tax=Endozoicomonas num...   452    6e-125
lcl|K2EN23  Uncharacterized protein n=1 Tax=uncultured bacterium ...   452    6e-125
lcl|G5JY56  Acetyl-CoA carboxylase, biotin carboxylase subunit n=...   451    6e-125
lcl|H8FRU9  Acetyl CoA carboxylase, biotin carboxylase subunit n=...   451    6e-125
lcl|K6X6Z8  Acetyl-CoA carboxylase, biotin carboxylase subunit n=...   451    7e-125
lcl|A0A085EF85  Acetyl-CoA carboxylase n=1 Tax=Flavobacterium sp....   451    8e-125
lcl|N6VT06  Methylcrotonoyl-coenzyme A carboxylase 1 (Alpha) n=1 ...   451    8e-125
lcl|U5F612  Acetyl-CoA carboxylase, biotin carboxylase subunit n=...   451    1e-124
lcl|L0HC90  Acetyl-CoA carboxylase, biotin carboxylase subunit n=...   451    1e-124
lcl|A0A073IXQ5  Acetyl-CoA carboxylase biotin carboxylase subunit...   451    1e-124
lcl|X8K5D3  Acetyl-CoA carboxylase, biotin carboxylase subunit n=...   450    2e-124
lcl|Q2BIU7  Biotin carboxylase n=2 Tax=Gammaproteobacteria RepID=...   450    2e-124
lcl|A8URI1  Pyruvate carboxylase n-terminal domain n=1 Tax=Hydrog...   450    2e-124
lcl|I2JPL9  Acetyl-CoA carboxylase, biotin carboxylase n=1 Tax=ga...   449    2e-124
lcl|S0L120  Acetyl-CoA carboxylase, biotin carboxylase subunit n=...   449    2e-124
lcl|G5KCU2  Acetyl-CoA carboxylase, biotin carboxylase subunit n=...   449    3e-124
lcl|A0A023BQE3  Biotin carboxylase n=4 Tax=Aquimarina RepID=A0A02...   449    3e-124
lcl|M5DH98  Biotin carboxylase of acetyl-CoA carboxylase n=1 Tax=...   449    3e-124
lcl|A0A062GRW7  Acetyl-CoA carboxylase, biotin carboxylase subuni...   449    3e-124
lcl|K7JAC5  Uncharacterized protein n=2 Tax=Nasonia vitripennis R...   449    3e-124
lcl|E3GX30  Pyruvate carboxylase subunit A n=1 Tax=Methanothermus...   449    4e-124
lcl|W9BDF8  AccC2 protein n=21 Tax=Enterobacteriaceae RepID=W9BDF...   449    5e-124
lcl|A0A073J124  Acetyl-CoA carboxylase n=11 Tax=Rhodobacteraceae ...   448    5e-124
lcl|E1RHS7  Acetyl-CoA carboxylase, biotin carboxylase n=1 Tax=Me...   448    5e-124
lcl|U2XVX3  Uncharacterized protein n=1 Tax=alpha proteobacterium...   448    7e-124
lcl|D7N3D9  Acetyl-CoA carboxylase, biotin carboxylase subunit n=...   448    7e-124
lcl|L8JDY6  Biotin carboxylase of acetyl-CoA carboxylase n=2 Tax=...   447    9e-124
lcl|I3IPW1  Biotin carboxylase n=1 Tax=planctomycete KSU-1 RepID=...   447    1e-123
lcl|S9RQ19  Biotin carboxylase of acetyl-CoA carboxylase n=1 Tax=...   447    2e-123
lcl|A0A087JWU3  Acetyl-CoA carboxylase n=293 Tax=Vibrionaceae Rep...   447    2e-123
lcl|A0A094NXA5  Acetyl-CoA carboxylase n=2 Tax=Cobetia RepID=A0A0...   447    2e-123
lcl|A0A091B1X2  Acetyl-CoA carboxylase biotin carboxylase subunit...   446    3e-123
lcl|A0A085HFL2  Biotin carboxylase n=3 Tax=Enterobacteriaceae Rep...   446    3e-123
lcl|A0A011N8T1  Biotin carboxylase n=2 Tax=Candidatus Accumulibac...   446    3e-123
lcl|A0A091ATW1  Acetyl-CoA carboxylase biotin carboxylase subunit...   446    3e-123
lcl|A0A095XKZ8  Acetyl-CoA carboxylase n=1 Tax=Peptostreptococcus...   446    4e-123
lcl|A8PLX8  Acetyl-CoA carboxylase, biotin carboxylase n=1 Tax=Ri...   446    4e-123
lcl|S1RK91  Acetyl-CoA carboxylase, biotin carboxylase subunit n=...   445    5e-123
lcl|S3DZT2  Acetyl-CoA carboxylase, biotin carboxylase n=2 Tax=Ca...   445    5e-123
lcl|A0A061QFE6  Biotin carboxylase n=1 Tax=alpha proteobacterium ...   445    6e-123
lcl|A6ENG5  Acetyl-CoA carboxylase, biotin carboxylase n=1 Tax=un...   445    6e-123
lcl|R9PM04  Biotin carboxylase of acetyl-CoA carboxylase n=1 Tax=...   445    6e-123
lcl|Q1MXR6  Acetyl-CoA carboxylase n=1 Tax=Bermanella marisrubri ...   445    7e-123
lcl|B8KW06  Acetyl-CoA carboxylase, biotin carboxylase n=1 Tax=Lu...   445    7e-123
lcl|A0A078M958  Acetyl-CoA carboxylase biotin carboxylase subunit...   445    7e-123
lcl|G2EC93  Pyruvate carboxylase subunit A n=1 Tax=Bizionia argen...   445    7e-123
lcl|A6FD94  Biotin carboxylase (A subunit of acetyl-CoA carboxyla...   444    8e-123
lcl|M7YT21  Acetyl-CoA carboxylase, biotin carboxylase n=1 Tax=Me...   444    9e-123
lcl|H3NVD0  Acetyl-CoA carboxylase, biotin carboxylase subunit n=...   444    1e-122
lcl|A4BFT5  Acetyl-CoA carboxylase n=1 Tax=Reinekea blandensis ME...   444    1e-122
lcl|E3R1J4  Acetyl-CoA carboxylase, biotin carboxylase subunit n=...   444    1e-122
lcl|H4GIL0  Acetyl-CoA carboxylase, biotin carboxylase n=1 Tax=La...   444    2e-122
lcl|F5SZL3  Biotin carboxylase n=2 Tax=Methylophaga RepID=F5SZL3_...   444    2e-122
lcl|N6W6G5  Pyruvate carboxylase subunit A n=1 Tax=Marinobacter n...   443    2e-122
lcl|W5YH53  3-methylcrotonyl-CoA carboxylase subunit alpha n=1 Ta...   443    2e-122
lcl|A0Z0J8  Acetyl-CoA carboxylase n=1 Tax=marine gamma proteobac...   443    2e-122
lcl|M7TDC6  Acetyl-CoA carboxylase, biotin carboxylase subunit n=...   443    2e-122
lcl|A7I988  Acetyl-CoA carboxylase, biotin carboxylase n=1 Tax=Me...   443    2e-122
lcl|G2J8R7  Biotin carboxylase (Acetyl-CoA carboxylase subunit A)...   443    3e-122
lcl|L8MPT7  Uncharacterized protein n=1 Tax=Pseudomonas pseudoalc...   442    3e-122
lcl|S2LFP4  Acetyl-CoA carboxylase biotin carboxylase subunit n=3...   442    4e-122
lcl|S7XRA3  Geranyl-CoA carboxylase biotin-containing subunit n=1...   441    8e-122
lcl|I9BS34  Pyruvate carboxylase n=1 Tax=Ralstonia sp. PBA RepID=...   441    9e-122
lcl|A3JUE7  Acetyl-CoA carboxylase n=1 Tax=Rhodobacteraceae bacte...   441    1e-121
lcl|V2UT28  Acetyl-CoA carboxylase, biotin carboxylase subunit n=...   441    1e-121
lcl|A0A095BQ63  Acetyl-CoA carboxylase, biotin carboxylase subuni...   441    1e-121
lcl|I2JKC4  Pyruvate carboxylase subunit A n=3 Tax=Gammaproteobac...   440    1e-121
lcl|E0E2H5  Acetyl-CoA carboxylase, biotin carboxylase subunit n=...   440    2e-121
lcl|W6LS98  Acetyl CoA carboxylase, biotin carboxylase subunit n=...   439    3e-121
lcl|R2P1W4  Acetyl-CoA carboxylase, biotin carboxylase subunit n=...   439    3e-121
lcl|T0JUS8  Biotin carboxylase 1 n=1 Tax=Sporomusa ovata DSM 2662...   439    3e-121
lcl|R2PXL4  Acetyl-CoA carboxylase, biotin carboxylase subunit n=...   439    3e-121
lcl|R3X0N9  Acetyl-CoA carboxylase, biotin carboxylase subunit n=...   439    3e-121
lcl|Q1N580  Acetyl/propionyl-CoA carboxylase, alpha subunit n=1 T...   439    4e-121
lcl|A0A076Z4Z1  Acetyl-CoA carboxylase n=230 Tax=Streptococcus ag...   439    4e-121
lcl|A0A090EV57  Propionyl-CoA carboxylase alpha chain, mitochondr...   439    4e-121
lcl|I3D5X0  Acetyl-CoA carboxylase, biotin carboxylase subunit n=...   439    4e-121
lcl|B8GGV2  Carbamoyl-phosphate synthase L chain ATP-binding n=1 ...   439    5e-121
lcl|B7RVT3  Acetyl-CoA carboxylase, biotin carboxylase n=1 Tax=ma...   439    5e-121
lcl|R8B5G9  Methylcrotonoyl-coenzyme A carboxylase 1 (Alpha) n=1 ...   439    5e-121
lcl|A0A074K0Q5  Acetyl-CoA carboxylase n=3 Tax=Thioclava RepID=A0...   439    5e-121
lcl|B6AYX3  Acetyl-CoA carboxylase, biotin carboxylase n=1 Tax=Rh...   438    6e-121
lcl|A0A094LRX7  Pyruvate carboxylase subunit A n=1 Tax=Shewanella...   438    6e-121
lcl|V7EHY9  Acetyl-CoA carboxylase n=1 Tax=Rhodobacter sp. CACIA1...   438    6e-121
lcl|M5FJP3  Propionyl-CoA carboxylase alpha chain,mitochondrial n...   438    7e-121
lcl|A0A010RJ13  3-methylcrotonyl-CoA carboxylase subunit alpha n=...   438    7e-121
lcl|N6WLR9  Pyruvate carboxylase subunit A n=1 Tax=Thermoplasmata...   438    7e-121
lcl|A0A095CSU7  Acetyl-CoA carboxylase n=1 Tax=Rhodovulum sp. NI2...   438    8e-121
lcl|K2QAE3  Propionyl-CoA carboxylase subunit alpha n=2 Tax=Rhizo...   438    9e-121
lcl|A9RNK1  Predicted protein n=1 Tax=Physcomitrella patens subsp...   437    1e-120
lcl|E8LL02  Acetyl-CoA carboxylase, biotin carboxylase subunit n=...   437    1e-120
lcl|K2BNB2  Uncharacterized protein n=1 Tax=uncultured bacterium ...   437    1e-120
lcl|S2DUE9  Methylcrotonyl-CoA carboxylase biotin-containing subu...   437    1e-120
lcl|A6F507  Acetyl-CoA carboxylase, biotin carboxylase n=1 Tax=Ma...   437    1e-120
lcl|N9MZG9  Acetyl-CoA carboxylase, biotin carboxylase subunit n=...   437    1e-120
lcl|A0A059IQB0  Acetyl-CoA carboxylase n=1 Tax=Defluviimonas sp. ...   437    2e-120
lcl|S9QJ56  Propionyl-CoA carboxylase biotin-containing subunit n...   437    2e-120
lcl|W6KLM9  Putative acyl-CoA carboxylase biotin-carrying subunit...   437    2e-120
lcl|W6TU52  Acetyl-CoA carboxylase n=1 Tax=Pedobacter sp. V48 Rep...   437    2e-120
lcl|A0A085L3F6  Acetyl-CoA carboxylase n=1 Tax=Schleiferia thermo...   437    2e-120
lcl|L8GNZ7  Carbamoylphosphate synthase L chain, ATP-binding, put...   437    2e-120
lcl|S7VR93  Pyruvate carboxylase subunit A n=1 Tax=Winogradskyell...   436    2e-120
lcl|U3H256  3-methylcrotonyl-CoA carboxylase subunit alpha n=1 Ta...   436    2e-120
lcl|S2LFL6  Acetyl-CoA carboxylase subunit alpha n=1 Tax=Halomona...   436    2e-120
lcl|A0A026WX95  Transportin-1 n=1 Tax=Cerapachys biroi RepID=A0A0...   436    3e-120
lcl|H1YXX6  Pyruvate carboxylase subunit A n=1 Tax=Methanoplanus ...   436    3e-120
lcl|A0A095V6U1  Acetyl-CoA carboxylase n=1 Tax=Rhizobium sp. YS-1...   436    3e-120
lcl|A1ZID5  Pyruvate carboxylase subunit A n=1 Tax=Microscilla ma...   436    4e-120
lcl|R1GQ60  Biotin carboxylase of acetyl-CoA carboxylase n=2 Tax=...   436    4e-120
lcl|U7M2Y2  Acetyl-CoA carboxylase, biotin carboxylase subunit n=...   435    5e-120
lcl|A0A073IHU9  Acetyl-CoA carboxylase n=4 Tax=Sulfitobacter RepI...   435    5e-120
lcl|A0A077GNK4  3-methylcrotonyl-CoA carboxylase n=699 Tax=Moraxe...   435    6e-120
lcl|A0A076MYV0  Acetyl/propionyl-CoA carboxylase, biotin carboxyl...   435    6e-120
lcl|A0A086XXG3  Acetyl-CoA carboxylase n=2 Tax=Haematobacter RepI...   435    6e-120
lcl|A0A074TGJ2  Acetyl-CoA carboxylase n=1 Tax=Thioclava dalianen...   435    7e-120
lcl|J2IFP7  Carbamoyl-phosphate synthase subunit L n=3 Tax=Alishe...   435    8e-120
lcl|J1IRW1  Geranyl-CoA carboxylase, alpha subunit AtuF n=4 Tax=P...   434    9e-120
lcl|A0A058ZLI1  Propionyl-CoA carboxylase subunit alpha n=1 Tax=A...   434    9e-120
lcl|S6HJ11  Biotin carboxylase n=2 Tax=unclassified Oceanospirill...   434    9e-120
lcl|A0A074MMI0  3-methylcrotonyl-CoA carboxylase n=1 Tax=Erythrob...   434    9e-120
lcl|A0A078MB96  Pyruvate carboxylase subunit A n=3 Tax=Pseudomona...   434    1e-119
lcl|J3EA85  Acetyl/propionyl-CoA carboxylase, alpha subunit n=3 T...   434    1e-119
lcl|A0P6Z1  Acetyl-CoA carboxylase n=1 Tax=Methylophilales bacter...   434    1e-119
lcl|A0A072NLA3  Acetyl-CoA carboxylase, biotin carboxylase subuni...   434    2e-119
lcl|A4EVF7  Propionyl-CoA carboxylase, alpha subunit n=2 Tax=Rhod...   433    2e-119
lcl|W1MTH4  3-methylcrotonyl-CoA carboxylase subunit alpha n=2 Ta...   433    2e-119
lcl|I2QJH6  Acetyl/propionyl-CoA carboxylase, alpha subunit n=2 T...   433    3e-119
lcl|M7NWJ7  Acetyl-/propionyl-coenzyme A carboxylase alpha chain ...   433    3e-119
lcl|A0A069PIB6  Acetyl-CoA carboxylase n=1 Tax=Burkholderia glath...   432    3e-119
lcl|G1WL24  Acetyl-CoA carboxylase n=1 Tax=Collinsella tanakaei Y...   432    3e-119
lcl|E2AJ02  Methylcrotonoyl-CoA carboxylase subunit alpha, mitoch...   432    4e-119
lcl|U2WUB7  Uncharacterized protein n=2 Tax=unclassified Alphapro...   432    4e-119
lcl|S5TLI2  Biotin carboxylase n=1 Tax=uncultured bacterium esnap...   432    5e-119
lcl|U3D4V2  Methylcrotonoyl-CoA carboxylase subunit alpha, mitoch...   432    5e-119
lcl|A0A081FT40  Pyruvate carboxylase subunit A n=3 Tax=Marinobact...   432    5e-119
lcl|B9SFG9  Acetyl-CoA carboxylase, putative n=1 Tax=Ricinus comm...   432    6e-119
lcl|K0B4U5  Carbamoyl-phosphate synthase L chain ATP-binding prot...   432    7e-119
lcl|D2U2K8  Biotin carboxylase n=2 Tax=Arsenophonus RepID=D2U2K8_...   431    7e-119
lcl|A0A023CWS3  Biotin carboxylase of acetyl-CoA carboxylase n=1 ...   431    7e-119
lcl|F7RVD4  Acetyl/propionyl-CoA carboxylase, alpha subunit n=1 T...   431    8e-119
lcl|W8I6H4  Propionyl-CoA carboxylase alpha chain protein n=3 Tax...   431    8e-119
lcl|H1S9G3  Methylcrotonoyl-CoA carboxylase n=1 Tax=Cupriavidus b...   431    9e-119
lcl|A0A069CWU0  Biotin carboxylase n=1 Tax=Weissella oryzae SG25 ...   431    1e-118
lcl|N7VAB4  Acetyl-CoA carboxylase, biotin carboxylase subunit n=...   431    1e-118
lcl|O30019  Pyruvate carboxylase subunit A n=2 Tax=Archaeoglobus ...   431    1e-118
lcl|K2HD97  Propionyl-CoA carboxylase, alpha subunit n=1 Tax=Ocea...   431    1e-118
lcl|A0A090DCQ5  Acetyl-CoA carboxylase, biotin carboxylase subuni...   431    1e-118
lcl|F7CQH6  DCN1-like protein n=10 Tax=Theria RepID=F7CQH6_MONDO       431    1e-118
lcl|A0A037ZLI6  Acetyl-CoA carboxylase n=1 Tax=Actibacterium muco...   431    1e-118
lcl|N9K8N5  Acetyl-CoA carboxylase, biotin carboxylase subunit n=...   431    1e-118
lcl|D7TE06  Putative uncharacterized protein n=1 Tax=Vitis vinife...   430    2e-118
lcl|H0T9W0  Putative biotin carboxylase n=1 Tax=Bradyrhizobium sp...   430    2e-118
lcl|H3NCQ0  Acetyl-CoA carboxylase, biotin carboxylase subunit n=...   430    2e-118
lcl|E1BGC1  Uncharacterized protein n=6 Tax=Cetartiodactyla RepID...   430    2e-118
lcl|A0A085XQR5  Biotin carboxylase of acetyl-CoA carboxylase n=1 ...   430    2e-118
lcl|W2G8T3  Acetyl-CoA carboxylase, biotin carboxylase subunit n=...   430    2e-118
lcl|S6K8X0  Biotin carboxylase/biotin-containing subunit n=1 Tax=...   430    2e-118
lcl|V6AUS9  Biotin carboxylase 1 n=1 Tax=Thaumarchaeota archaeon ...   430    2e-118
lcl|I1AQB8  Propionyl-CoA carboxylase, alpha subunit n=1 Tax=Citr...   430    2e-118
lcl|Q4PKB0  Predicted acetyl-CoA carboxylase n=2 Tax=Bacteria Rep...   429    3e-118
lcl|A3TYU3  Propionyl-CoA carboxylase, alpha subunit n=1 Tax=Ocea...   429    3e-118
lcl|Q2QMG2  Methylcrotonoyl-CoA carboxylase subunit alpha, mitoch...   429    4e-118
lcl|D0N1Q9  Methylcrotonoyl-CoA carboxylase subunit alpha, putati...   429    4e-118
lcl|A0A090LBC8  Uncharacterized protein n=1 Tax=Strongyloides rat...   429    4e-118
lcl|A2SPQ9  Pyruvate carboxylase subunit A n=1 Tax=Methanocorpusc...   429    4e-118
lcl|B6B1N1  RimK-like ATP-grasp domain family n=1 Tax=Rhodobacter...   429    5e-118
lcl|R2NF54  Acetyl-CoA carboxylase, biotin carboxylase subunit n=...   429    5e-118
lcl|W8ANS0  Methylcrotonyl-CoA carboxylase biotin-containing subu...   429    5e-118
lcl|Q99MR8  Methylcrotonoyl-CoA carboxylase subunit alpha, mitoch...   429    5e-118
lcl|W5G176  Uncharacterized protein n=6 Tax=Triticeae RepID=W5G17...   428    6e-118
lcl|I4YKN7  Acetyl/propionyl-CoA carboxylase, alpha subunit n=2 T...   428    6e-118
lcl|F7NW79  Acetyl/propionyl-CoA carboxylase, alpha subunit n=1 T...   428    7e-118
lcl|M5J509  Acetyl-CoA carboxylase, biotin carboxylase n=2 Tax=La...   428    7e-118
lcl|R0ERG8  Acetyl-CoA carboxylase, biotin carboxylase subunit, a...   428    8e-118
lcl|V4ASR2  Uncharacterized protein n=1 Tax=Lottia gigantea RepID...   428    8e-118
lcl|K2PIS1  Carbamoyl-phosphate synthase subunit L n=1 Tax=Nitrat...   428    8e-118
lcl|A0A086Y2A7  3-methylcrotonyl-CoA carboxylase n=1 Tax=Haematob...   428    8e-118
lcl|N8YSH7  Acetyl-CoA carboxylase, biotin carboxylase subunit n=...   428    9e-118
lcl|E3C9Z5  Putative acetyl-CoA carboxylase, biotin carboxylase s...   428    9e-118
lcl|A0A086D2N7  Pyruvate carboxylase subunit A n=1 Tax=Gammaprote...   428    9e-118
lcl|W4HUM1  Biotin carboxyl carrier protein n=1 Tax=Mycobacterium...   427    1e-117
lcl|A7RUP0  Predicted protein n=3 Tax=Nematostella vectensis RepI...   427    1e-117
lcl|H1LD79  Putative acetyl-CoA carboxylase, biotin carboxylase s...   427    1e-117
lcl|E0MIY6  Propionyl-CoA carboxylase alpha chain n=1 Tax=Ahrensi...   427    1e-117
lcl|Q0FU36  Propionyl-CoA carboxylase, alpha subunit n=2 Tax=Rhod...   427    1e-117
lcl|A0A081M800  Acetyl-CoA carboxylase n=3 Tax=Rhizobium RepID=A0...   427    1e-117
```
