## Supplemental Table 3 for "Oligomeric assemblies of plant biotin carboxylase revealed by cryo-EM and cross-linking"

Table S3. Evolutionary importance of each sequence position of *Brassica* BC. Lower coverage and lower score implies higher functional or structural importance.

```
%	 Column details:
%
% alignment# is the alignment position in the MSA.
% residue# is the residue position in the query protein.
% type is the one letter amino acid name in the query protein.
% coverage is the relative importance position for each residue. Low coverage implies evolutionary importance.
% variability shows the residue variation in the column of the MSA.
% Evolutionary scores for residues are calculated with rvET method. Low score implies evolutionary importance.
% Mihalek I, Res I, Lichtarge O. (2004). A Family of Evolution-Entropy Hybrid Methods for Ranking of Protein Residues by Importance. J. Mol. Bio. 336(5): 1265-82.
%
%	 RESIDUE RANKS:
% alignment#  residue#   AA type  coverage              variability       Score(rvET)
         1       74          K   0.25727    6                     KTMR.S     11.48
         2       75          I   0.69128    5                      VIL.M     52.88
         3       76          L   0.24609    6                     LMFVI.     10.95
         4       77          V   0.48546    4                       IV.F     29.95
         5       78          A   0.20805    4                       ASV.      7.82
         6       79          N   0.13199    2                         N.      3.08
         7       80          R   0.13199    2                         R.      3.08
         8       81          G   0.13199    2                         G.      3.08
         9       82          E   0.13647    3                        ED.      3.26
        10       83          I   0.14541    3                        IV.      3.68
        11       84          A   0.17002    4                       AVS.      4.89
        12       85          V   0.39374    8                   LIVCTAM.     22.26
        13       86          R   0.15884    3                        RQ.      4.26
        14       87          V   0.58837    4                       VIA.     39.56
        15       88          I   0.58613   11                IMLQTAVEHF.     39.56
        16       89          R   0.38702    9                  RKTQEHDA.     22.09
        17       90          T   0.37584    7                    TSAEVG.     21.42
        18       91          A   0.30201    5                      CALI.     15.40
        19       92          H   0.72483   12               NKQRTHALIW.E     58.64
        20       93          E   0.51454   10                 AKEQRTNDS.     32.87
        21       94          M   0.55481    7                    LMKNAQ.     36.46
        22       95          G   0.38031    8                   GNEDKSQ.     21.48
        23       96          I   0.41834    8                   ICHLYVKM     24.60
        24       97          P   0.87472   15            KRTAGSEQPLHVDNI     81.66
        25       98          C   0.46532    7                    TSVCAPM     28.39
        26       99          V   0.23490    4                       VAIL      9.78
        27      100          A   0.21253    7                    AQLVIMG      8.03
        28      101          V   0.50112    6                     VIAGTL     31.67
        29      102          Y   0.40045    6                     YFHCVW     22.47
        30      103          S   0.19016    4                       SNTA      6.06
        31      104          T   0.67785   11                DETVSNKQLAM     51.70
        32      105          I   0.47204    7                    EAVPGIS     29.10
        33      106          D   0.11857    3                        DEK      2.85
        34      107          K   0.78523   14             YRAKHSVEQTDNC.     70.44
        35      108          D   0.87696   16           NGEQHDRKATYFSMP.     81.70
        36      109          A   0.51230    7                    SAGTLC.     32.80
        37      110          L   0.49441   15            LPTVMREAKQSHFY.     31.44
        38      111          H   0.24385    6                     HAYPF.     10.69
        39      112          V   0.38479    9                  VARLTPIS.     21.95
        40      113          K   0.77405   17          KRLQDEASHTYFNVIM.     67.92
        41      114          L   0.65101   16           KFMYAEQLDTVSRIH.     47.88
        42      115          A   0.14318    4                       ASV.      3.52
        43      116          D   0.21029    7                    DGETNH.      7.84
        44      117          E   0.34676   11                EQTRIAMHYS.     19.75
        45      118          A   0.58389   10                 SARGTVKCD.     39.38
        46      119          V   0.50559    8                   YVIFHWL.     32.22
        47      120          C   0.61074   16           HLCRWYNFSMQPAEV.     43.64
        48      121          I   0.46756    5                      ILVA.     28.49
        49      122          G   0.15436    5                      GSNV.      4.02
        50      123          E   0.61969   12               EKPGAVDSNQT.     44.31
        51      124          A   0.71365   14             APSGDNEHQTVIK.     57.04
        52      125          P   0.83893   16           APTLQKSMDNERHYI.     76.47
        53      126          S   0.59060    9                  PSTAVL.IG     39.64
        54      127          N   0.88591   15            AKNSRGTQLDEPIVM     83.90
        55      128          Q   0.82327   12               KEQALDSTRGNM     74.84
        56      129          S   0.13870    4                       STGE      3.35
        57      130          Y   0.10067    2                         YS      2.10
        58      131          L   0.18345    5                      LINGT      5.89
        59      132          L   0.62192   13              NVLRKQIDSCAHE     44.46
        60      133          I   0.79195   14             QMGAISVLTYKPFE     70.81
        61      134          P   0.79642   17          EDSGAPTRHQLIYNVFK     71.55
        62      135          N   0.64430   12               KTRANSHGQLME     47.02
        63      136          V   0.44743    4                       IVLM     26.42
        64      137          L   0.73602    5                      LMIVQ     59.27
        65      138          S   0.74944   11                EQKADTNSYGH     62.81
        66      139          A   0.45190    7                    VAITLNS     26.67
        67      140          A   0.37360    8                   MIASCVTF     21.41
        68      141          I   0.76063   15            LKRTAVHQIEMSDNC     65.25
        69      142          S   0.82550   17          SADETQVKRHILGNMF.     75.26
        70      143          R   0.63758   15            STLICAVMFYGKRH.     45.44
        71      144          G   0.57718   11                GNQHKARDES.     39.12
        72      145          C   0.59732    7                    TAVSCI.     41.28
        73      146          T   0.60850   12               DEQNKVTSMIA.     43.60
        74      147          M   0.29306    7                    AGCSL.M     14.96
        75      148          L   0.47427    4                       VIL.     29.21
        76      149          H   0.11633    4                       HFY.      2.75
        77      150          P   0.10067    2                         P.      2.10
        78      151          G   0.10067    2                         G.      2.10
        79      152          Y   0.21477    5                      YFVI.      8.41
        80      153          G   0.10067    2                         G.      2.10
        81      154          F   0.10067    2                         F.      2.10
        82      155          L   0.15213    4                       LFM.      4.00
        83      156          A   0.34899    3                        SA.     19.93
        84      157          E   0.10067    2                         E.      2.10
        85      158          N   0.22148    7                    NRSDVK.      8.94
        86      159          A   0.72260   15            DSAEKPMTQYVLHG.     57.95
        87      160          L   0.92841   15            DVNEGTSALRKHQYP     92.43
        88      161          F   0.19687    3                        FLW      6.53
        89      162          V   0.35347    7                    APCSVGN     20.08
        90      163          E   0.81655   12               RKLEQADSNTM.     74.51
        91      164          M   0.74049   17          LMAKSERTIVNDCQYF.     60.43
        92      165          C   0.40268    6                     CAVLI.     22.57
        93      166          R   0.84787   16           ETNKSADRQVHIGLM.     77.78
        94      167          D   0.90828   14             KDAERNQSVHMTG.     88.84
        95      168          H   0.70470   16           NASEMTHQKRYCVLF.     54.43
        96      169          R   0.65324   11                KGDNSREHAQ.     48.26
        97      170          I   0.67338    9                  ILVFMYQA.     50.81
        98      171          N   0.89262   17          TNIVQKAERCGSLHDM.     85.02
        99      172          F   0.21700    4                       FWL.      8.45
       100      173          I   0.41163    4                       IVL.     23.91
       101      174          G   0.10067    2                         G.      2.10
       102      175          P   0.12528    3                        PA.      3.04
       103      176          N   0.86353   15            SKDGRTPNHAEQFL.     78.87
       104      177          P   0.82103   16           ASPTVKWDGLHYEIF.     74.55
       105      178          D   0.97315   16           DEQHKSGNVPRATIY.    100.12
       106      179          S   0.61745   12               SAVITQMHLNC.     44.16
       107      180          I   0.24832    4                       MIL.     11.00
       108      181          R   0.82774   16           NDERKIAQVSHTGLY.     75.53
       109      182          V   0.86577   17          LVIASKDTGRMQENFY.     79.24
       110      183          M   0.21924    7                    CMLVAI.      8.76
       111      184          G   0.14989    4                       GAS.      3.84
       112      185          D   0.42506    8                   DNSILHTA     25.23
       113      186          K   0.06264    1                          K      1.00
       114      187          S   0.57047   13              MLIARSNTVDEQG     38.26
       115      188          T   0.76734   15            ERASTGLVQHKNMDI     66.22
       116      189          A   0.22819    4                       CASL      9.36
       117      190          R   0.30425    4                       KRIG     15.62
       118      191          E   0.91723   16           AEDNKRTILSQFYHVM     89.72
       119      192          T   0.69351   14             ATLIREVSMFCHKQ     53.43
       120      193          M   0.25951    4                       MVAC     11.52
       121      194          K   0.90157   15            LNKSAMQIGTVREDH     87.19
       122      195          N   0.98434   12               KSDNHAQERTGF    103.62
       123      196          A   0.44295   14             AYTFCSLNVQIHGR     26.26
       124      197          G   0.57271    9                  KNGDRQSHE     38.29
       125      198          V   0.27517    5                      VILMC     13.53
       126      199          P   0.16555    5                      PNSQH      4.74
       127      200          T   0.57942    6                     TMLCVI     39.22
       128      201          V   0.52125    7                    VITLAMS     33.21
       129      202          P   0.27069    9                  PKGNAEQLM     13.14
       130      203          G   0.10738    3                        GWY      2.36
       131      204          S   0.50783   12               SILTVFHYARDG     32.51
       132      205          D   0.79418   16           PEDALMQHV.KSGNTR     71.37
       133      206          G   0.55705   12               GDEHKRQA.STN     37.01
       134      207          L   0.81208   18         LAEVPRDSNTKIGFMY.Q     74.04
       135      208          L   0.69575   14             VILGDNQREHSKM.     53.82
       136      209          Q   0.91051   18         DKTSNEAGLQHR.VPMYF     89.09
       138      210          S   0.86130   11                TDSGNREPVIH     78.82
       139      211          T   0.94855   21      AVIDPSTENFLMRQYHK.GWC     94.90
       140      212          E   0.93512   14             DEKAPQGTSVHIRN     93.91
       141      213          E   0.89038   19        ELKAQHDVRTYCFSI.NMG     84.67
       142      214          G   0.50336   12               AGVLMFICNTKS     31.83
       143      215          V   0.93960   16           ELKQVIACFRSDHYMG     94.19
       144      216          R   0.99776   15            KDQLTIRVEASHNGM    119.45
       145      217          L   0.83445   15            IVRKAEQHLFMCTNW     76.12
       146      218          A   0.33557    9                  ASGLCVIEN     18.57
       147      219          N   0.98658   14             NKSVQGDTRAEHFL    105.17
       148      220          E   0.97763   15            EQDSGRKAHTLNVIY    103.07
       149      221          I   0.56823    7                    IVMLTAF     37.76
       150      222          G   0.26398    8                   GKSMAHTE     12.31
       151      223          F   0.48098    4                       YFWL     29.80
       152      224          P   0.06264    1                          P      1.00
       153      225          V   0.46085    6                     VILAQC     28.07
       154      226          M   0.42953    5                      LMIVA     25.50
       155      227          I   0.55034    4                       LIVM     36.08
       156      228          K   0.06264    1                          K      1.00
       157      229          A   0.17226    3                        SAP      5.02
       158      230          T   0.48322    6                     VSATRC     29.83
       159      231          A   0.43848   11                YARQHLSNTGD     26.24
       160      232          G   0.06264    1                          G      1.00
       161      233          G   0.06264    1                          G      1.00
       162      234          G   0.06264    1                          G      1.00
       163      235          G   0.06264    1                          G      1.00
       164      236          R   0.29978    5                      RKIMV     15.27
       165      237          G   0.06488    2                         GS      1.04
       166      238          M   0.20582    2                         IM      7.64
       167      239          R   0.25280   11                RFKQSVTHNAG     11.22
       168      240          L   0.60403   12               LVIARQMPTKEF     42.80
       169      241          A   0.40492    5                      VAICS     22.96
       170      242          N   0.94183   17          TNEDWYSAPQKRHLMFG     94.23
       171      243          E   0.95749   12               NKQESDTGARCH     95.98
       172      244          P   0.80313   15            DLEAPMSKVGTRIQN     72.04
       173      245          S   0.97539   16           KQEDASGNRTPVILCH    102.93
       174      246          E   0.83221   14             EDGRNSAQTKLMHV     75.98
       175      247          F   0.54810    8                   LFACVIMT     35.70
       176      248          V   0.96868   16           QRPELAKVIHDMSYTG     98.05
       177      249          K   0.91946   16           ESDAQPKRHTNGVFML     90.68
       178      250          L   0.73378   15            GAQLENHMSYVDKTC     59.19
       179      251          L   0.70022    8                   FYMLIVWA     54.19
       180      252          Q   0.98881   19        EDQKNRAHVLFGPTWSMYI    106.11
       181      253          Q   0.72707   16           TRLSGANEQ.IVMFKD     58.69
       182      254          A   0.32886    8                   VASCT.IG     17.91
       183      255          K   0.67114   13              TIVKRQYS.EMGA     50.75
       184      256          S   0.71812   14             SGANRT.KELQMIH     57.74
       185      257          E   0.19911    9                  EN.DLTIVM      6.68
       186      258          A   0.27293    6                     SAG.VT     13.33
       187      259          A   0.93065   15            IKTVEAGLQHMSRND     93.07
       188      260          A   0.76286   14             SANGQHKVRMILTE     65.48
       189      261          A   0.55257   13              ASGLDFCNWVYTI     36.31
       190      262          F   0.18568    4                       VFLY      5.94
       191      263          G   0.38255    7                    GNSTKDR     21.58
       192      264          N   0.43624   10                 KDSINCRTEV     26.11
       193      265          D   0.75168   11                SGDAPQNTEKR     63.49
       194      266          G   0.68233   14             ASRTEIKDVGQMNH     51.72
       195      267          V   0.56376    6                     IVMLFC     37.53
       196      268          Y   0.26846    4                       ILFY     12.87
       197      269          L   0.70246    6                     VILMAC     54.43
       198      270          E   0.06264    1                          E      1.00
       199      271          K   0.29530    4                       KRQH     15.06
       200      272          Y   0.53468    8                   FYACKILV     34.82
       201      273          V   0.53915    6                     LVIMFC     35.26
       202      274          Q   0.76510   18         EGSATDLIQVMRKNFPHY     66.07
       203      275          N   0.79866   16           KSNEDQRAYTHPLIGC     71.62
       204      276          P   0.26174    8                   TPSAIVFM     11.64
       205      277          R   0.28859    3                        RKH     14.87
       206      278          H   0.06264    1                          H      1.00
       207      279          I   0.45414    4                       IVLC     26.68
       208      280          E   0.06264    1                          E      1.00
       209      281          F   0.53244    7                    YIVMLFA     34.59
       210      282          Q   0.06264    1                          Q      1.00
       211      283          I   0.63311    4                       MVIL     45.35
       212      284          L   0.60626    7                    CALMIFV     43.58
       213      285          A   0.51902    6                     RAGCSV     33.20
       214      286          D   0.07383    2                         DN      1.36
       215      287          K   0.96421   14             HKTSMQAGRNELDC     96.66
       216      288          F   0.85906   14             HQFDYEKLNRAMST     78.64
       217      289          G   0.25056    8                   GE.NRKQD     11.08
       218      290          N   0.65772   10                 NHDTGRKESQ     49.05
       219      291          V   0.75615   12               ATIVCLGYSRMH     64.66
       220      292          V   0.71141    5                      VILAC     56.97
       221      293          H   0.42729    9                  HYWAFSQTD     25.36
       222      294          F   0.28188    7                    LVIFCMA     14.45
       223      295          G   0.53691   12               FNGHYACWLPSM     35.20
       224      296          E   0.22595    4                       EADT      9.34
       225      297          R   0.06264    1                          R      1.00
       226      298          D   0.20358    3                        EDN      7.64
       227      299          C   0.06264    1                          C      1.00
       228      300          S   0.17673    2                         ST      5.07
       229      301          I   0.60179    9                  IVLAMTSCF     42.07
       230      302          Q   0.06264    1                          Q      1.00
       231      303          R   0.12081    3                        RMI      2.85
       232      304          R   0.22371    5                      RNKTS      9.02
       233      305          N   0.37808    9                  NHQRSYKML     21.42
       234      306          Q   0.06264    1                          Q      1.00
       235      307          K   0.06264    1                          K      1.00
       236      308          L   0.41611    4                       LVIM     24.42
       237      309          L   0.63087    8                   IVLWFMAT     44.82
       238      310          E   0.06264    1                          E      1.00
       239      311          E   0.14765    6                     QEIVLM      3.73
       240      312          A   0.35794    5                      TASCG     20.46
       241      313          P   0.16331    6                     PLGNTQ      4.66
       242      314          S   0.36689    6                     SCAGPD     20.82
       243      315          P   0.54139   16           PAVSLEFTMHKIRDCN     35.42
       246      316          A   0.72036   16           VILFAGNQYSCDHMTR     57.81
       247      317          L   0.49888    6                     VLMIFY     31.58
       248      318          T   0.75391   12               DTSGKNMVPRAQ     64.24
       249      319          P   0.97092   18         AQPESIDLTNKHWVRFYG     99.96
       250      320          E   0.89933   14             AKEDQSLNRGTHMV     86.11
       251      321          L   0.74273   14             KTIVLMFQEARSHN     61.70
       252      322          R   0.17897    8                   RATLIVCQ      5.48
       253      323          K   0.99553   15            EDKQNSRAGVTILHM    112.66
       254      324          A   0.92394   15            EMAKRLDSQHYTNFI     91.53
       255      325          M   0.45861    7                    IVMLYTA     28.04
       256      326          G   0.31096   13              GCYAHLQFTISNM     16.05
       257      327          D   0.94407   14             EDQAKTSGRNMLVH     94.60
       258      328          A   0.61521   14             LACQETVIMKSDRY     43.96
       259      329          A   0.32438    6                     VASCGT     17.24
       260      330          V   0.62864   10                 VICLATSRMK     44.60
       261      331          A   0.85459   15            KNDAQETRSVLHMIY     78.55
       262      332          A   0.28412    8                   AVLTGIMC     14.48
       263      333          A   0.46980    9                  ASVGTCLIM     28.54
       264      334          A   0.84564   12               EKQRALSGVIHN     77.06
       265      335          S   0.72931   17          ASNTEQGYHKRDVLCIF     59.16
       266      336          I   0.54586    9                  VCILTAMFS     35.51
       267      337          G   0.80761   10                 DNSGARKHQE     72.53
       268      338          Y   0.07159    2                         YF      1.34
       269      339          I   0.87025   16           TVLIYAEDQCHFRKSN     80.58
       270      340          G   0.34452    3                        NGS     19.67
       271      341          V   0.25503    6                     LAVTSM     11.33
       272      342          G   0.06264    1                          G      1.00
       273      343          T   0.08501    2                         TS      2.06
       274      344          V   0.42282    8                   AVLMIFYC     25.04
       275      345          E   0.06264    1                          E      1.00
       276      346          F   0.18792    3                        FYC      6.06
       277      347          L   0.26622    4                       LIVM     12.46
       278      348          L   0.61298   12               RLIVASMFCYWT     43.69
       279      349          D   0.40716   10                 ADGSTKNEQ.     23.25
       281      350          E   0.70694   15            DEQKRGASPTIHN.V     55.23
       282      351          R   0.97987   18         SNHQKEDARILVT.GYFM    103.16
       285      352          G   0.63535   14             GKLHRQDENV.YMS     45.42
       286      353          S   0.88814   12               ENKDSQRA.GHT     84.65
       288      354          F   0.35123    5                      FYLVI     19.99
       289      355          Y   0.32215    6                     YFWHCA     17.24
       290      356          F   0.06264    1                          F      1.00
       291      357          M   0.27740    5                      ILMTC     13.75
       292      358          E   0.06264    1                          E      1.00
       293      359          M   0.15660    4                       IMVA      4.10
       294      360          N   0.06264    1                          N      1.00
       295      361          T   0.11186    5                      ATCPS      2.53
       296      362          R   0.06264    1                          R      1.00
       297      363          I   0.36465    3                        LVI     20.80
       298      364          Q   0.06264    1                          Q      1.00
       299      365          V   0.11409    3                        VAI      2.65
       300      366          E   0.10291    3                        EKR      2.11
       301      367          H   0.07830    2                         HP      1.36
       302      368          P   0.29083    7                    PTACSGR     14.96
       303      369          V   0.33110    2                         IV     17.97
       304      370          T   0.31544    3                        STR     16.14
       305      371          E   0.07830    2                         EA      1.36
       306      372          M   0.74497   15            MLFWQKAETCSGVIY     61.79
       307      373          I   0.51678    5                      VILTA     33.12
       308      374          Y   0.35570   10                 STAVWCYFNI     20.11
       309      375          S   0.31767    6                     GKDSRN     16.50
       311      376          V   0.80089   11                LITVMKQYFCG     71.94
       312      377          D   0.12304    3                        DNE      3.01
       313      378          L   0.44072    6                     FLIVMT     26.24
       314      379          I   0.40940    5                      VIALT     23.28
       315      380          E   0.63982   13              KEAQRSNHIVCML     45.56
       316      381          E   0.59284   13              LQMEWCAKDVTHY     39.95
       317      382          Q   0.19463    3                        QMG      6.47
       318      383          I   0.57494    5                      ILVMF     38.60
       319      384          R   0.76957   16           DKRHLQAFCNSIMTYE     66.41
       320      385          V   0.68009    6                     IVLNMA     51.72
       321      386          A   0.18121    3                        ASG      5.59
       322      387          M   0.84116   17          NREAYSDQVGCHMLTFW     76.91
       323      388          G   0.24161    6                     GNDHSK     10.56
       324      389          E   0.78971   17          EKQHDRLAMYNGSVFIC     70.54
       325      390          K   0.90604   15            PKSTHAVIEQDMCRG     88.64
       326      391          L   0.30872    6                     LIFMVA     16.04
       327      392          R   0.88367   16           PANSTKRGDEQWHCVL     83.42
       333      393          Y   0.78300   13              FLIYAVMK.HWCR     70.18
       335      394          K   0.99329   16           KTSQRDGANPWEHLCF    108.42
       336      395          Q   0.19239    8                   QMWSDKRT      6.22
       337      396          E   0.91499   11                KEQDSAHNRGL     89.43
       338      397          E   0.62416   10                 DEGQAKSNYP     44.46
       339      398          I   0.70917   10                 LVIAWMPFTY     55.75
       340      399          V   1.00000   17          KRESQPTGHLVNADICM    122.66
       341      400          L   0.77852   18         MILFYHKRQNSACPVEWD     69.31
       342      401          R   0.95078   19        NKSQRTIADHMLEFVWG.Y     95.36
       343      402          G   0.17450    7                    GYKCRLM      5.04
       344      403          H   0.34004   12               YHWFAVQMSRNC     19.15
       345      404          S   0.36913    3                        ASV     21.25
       346      405          I   0.59508    6                     ILMVFA     40.63
       347      406          E   0.13423    2                         EQ      3.12
       348      407          C   0.36018   10                 CVLSNATFIY     20.67
       349      408          R   0.06264    1                          R      1.00
       350      409          I   0.33781    3                        IVL     18.62
       351      410          N   0.23043    5                      NYCAT      9.50
       352      411          A   0.14094    3                        ASC      3.47
       353      412          E   0.06935    2                         EY      1.32
       354      413          D   0.38926   10                 DANTHKRMIS     22.17
       355      414          P   0.41387   10                 TPAVSWGLIM     24.07
       356      415          F   0.80537   20       FLMEAKYVGRDSNQPT.IWC     72.05
       357      416          K   0.80984   20       LANDRQHTSGKIFEVPY.MC     73.64
       358      417          G   0.75839   12               DNGAQESTKRH.     65.24
       359      418          F   0.06711    2                         FL      1.31
       360      419          R   0.69799   13              ALMTRYQVSIFNK     54.05
       361      420          P   0.10962    2                         PA      2.53
       362      421          G   0.58166   13              SNDTQAGFCRHWM     39.31
       363      422          P   0.43177    9                  TIVGSAFPC     25.97
       364      423          G   0.08054    3                        GAS      1.52
       365      424          R   0.86801   16           PTNRHVKAEDQIMLCY     80.05
       366      425          I   0.43400    6                     VLMIFT     26.10
       367      426          T   0.87919   17          PTEDQSRVLAHFYKNGI     82.14
       368      427          S   0.87248   17          DTIRVHKQALSYENFGM     81.54
       369      428          Y   0.52573    9                  VYLWFIMCA     33.45
       370      429          L   0.64206   17          TKQRIHVASDEYFNLCM     45.88
       371      430          P   0.74720   15            ITLRKEPMVAYHQSF     62.02
       372      431          S   0.28635    8                   P.AKESVI     14.63
       374      432          G   0.64877   17          ASTKVEQH.DGLNPRMC     47.27
       379      433          G   0.36242   13              GAESPML.QVTDN     20.78
       401      434          P   0.83669   19        PEKNAGIVTHYDSQLMRFC     76.28
       402      435          F   0.53020   16           NGSYTVIAEHDCFKQW     34.48
       403      436          V   0.56152    6                     VILATM     37.42
       404      437          R   0.06264    1                          R      1.00
       405      438          M   0.68680   10                 CVLNTFMYIW     52.28
       406      439          D   0.23714    2                         DE      9.82
       407      440          S   0.49664    7                    TDNASHG     31.55
       408      441          H   0.47651   14             YGSAHLFQVPDCMI     29.23
       409      442          V   0.64653    8                   LFIVMASC     47.26
       410      443          Y   0.52796   16           YEKQTFLRGAVISCHN     33.66
       411      444          P   0.81432   15            PEQDASTRKCGNHVM     74.26
       412      445          D   0.39150   11                GQADNHKSMER     22.21
       413      446          Y   0.44966   15            CMGSTAEQDFYLIVW     26.66
       414      447          V   0.89709   18         TDEQVANMPRSIHLKCFY     86.00
       416      448          V   0.37136    4                       VILP     21.32
       417      449          P   0.45638    7                    SPGTQLV     26.69
       418      450          P   0.42058   12               PIVMLSTGCEAH     24.74
       419      451          S   0.66890   12               FYDHSINELMCT     50.72
       420      452          Y   0.16107    2                         YF      4.34
       421      453          D   0.06264    1                          D      1.00
       422      454          S   0.16779    3                        SPN      4.83
       423      455          L   0.30649    2                         LM     15.72
       424      456          L   0.47875    6                     MLIVCA     29.32
       425      457          G   0.49217    6                     ASGLVM     30.51
       426      458          K   0.06264    1                          K      1.00
       427      459          L   0.48770    5                      LVMIF     30.09
       428      460          I   0.39821    6                     CIVTSL     22.43
       429      461          V   0.55928    7                    TSVAGIC     37.17
       430      462          W   0.66443   13              WYHSRIMKLVFCT     50.46
       431      463          A   0.54362   10                 GDASQKLMRE     35.49
       432      464          P   0.93289   17          PQKEADRGSIMLNTHFV     93.18
       433      465          T   0.81879    9                  TDSNEGLHK     74.55
       440      466          R   0.10515    4                       FRWY      2.32
       441      467          E   0.98210   18         EDAKLIQGTSNPHVRMFC    103.33
       442      468          R   0.77181   16           EQKHTAMSDRIVGFLY     67.89
       443      469          A   0.34228    7                    SACLVTN     19.58
       444      470          I   0.56600    9                  RITVLCAMF     37.59
       445      471          E   0.95526   16           TLQEMKDARSGNVHIW     95.80
       446      472          R   0.48993   12               RLIKSHATGEVC     30.20
       447      473          M   0.39597   10                 MQLTSGAIVR     22.31
       448      474          K   0.78076   16           LVISQAGRDMKTNECH     70.07
       449      475          R   0.66667   18         TNKESRYDHVLACWQGMI     50.70
       450      476          A   0.32662    8                   AGKSCTVL     17.77
       451      477          L   0.23937    4                       LIVM     10.30
       452      478          N   0.88143   16           NDKAERQSHTGVWYFL     82.68
       453      479          D   0.46309   11                DNAETGRSQKM     28.25
       454      480          T   0.67562   10                 MFYTALCSIV     51.24
       455      481          I   0.78747   17          YNKEHIDQAVRTGSLFM     70.48
       456      482          I   0.68456    8                   VILAGFST     52.12
       457      483          T   0.77629   17          EQKTVRDLFASYHGNMI     68.03
       458      484          G   0.08277    3                        GPE      1.60
       461      485          V   0.59955    5                      VILPY     41.54
       462      486          P   0.85235   19        ESMQKANRGTPDHCVILYF     78.20
       463      487          T   0.31320    7                    THSGNCI     16.10
       464      488          T   0.23266    6                     STNILV      9.68
       465      489          I   0.65996   12               ILVTQRKEAMHS     49.05
       466      490          E   0.73154   16           PASGDEQVHYRNTILK     59.19
       467      491          Y   0.33333    6                     LFYIMV     18.19
       468      492          H   0.52349   16           YGCLVAIHFNQTSEDM     33.30
       469      493          K   0.85682   17          KTRASQLMGVYHIENDF     78.57
       470      494          L   0.85011   17          TFSARCNGDEKQMLWYV     77.85
       471      495          I   0.51007    7                    IVLACTM     32.64
       473      496          L   0.66219    9                  LCMFIASVT     49.56
       474      497          E   0.96644   18         NKEQDSARVGTHCLMFYI     97.42
       475      498          V   0.68904   14             SHNQDKETALGIRV     52.68
       476      499          E   0.90380   15            ENDTQPASKHWVGRL     87.47
       477      500          D   0.92170   19        EARKIDVGTSCHQLYNPFM     90.99
       478      501          F   0.31991    8                   YFWVILME     17.05
       479      502          K   0.92617   16           KTRIQVLAGECMNHSF     91.93
       480      503          N   0.99105   13              NRSEDATKQGHFL    107.55
       481      504          G   0.27964    9                  GRANSVQTD     14.22
       483      505          K   0.94631   15            EDNHQKRASLGTCIM     94.78
       484      506          V   0.62640   11                LFIAMTVHYCQ     44.54
       485      507          D   0.65548   11                SNDTHEGYRAF     48.64
       486      508          T   0.20134    5                      TINAV      7.32
       487      509          A   0.73826   14             DHNKGASRQCYT.E     60.10
       488      510          F   0.29754    9                  FYLWTVH.I     15.22
       489      511          I   0.44519    6                     LVIMFC     26.30
       490      512          P   0.71588   14             KSENPADGQRYTMI     57.07
       491      513          K   0.89485   15            RSKNEDAQHGTIVML     85.51
       492      514          H   0.84340   21      YHEQRIDANFWSLKMGPVTC.     77.02
       493      515          E   0.82998   21      GNYFLARCQMSHKPIEV.DTW     75.97
       494      516          E   0.91275   20       MSNDITPKAEVHGRQL.FWC     89.42
       495      517          E   0.95302   20       IPASEDQGWTVLFK.NMRHY     95.77
       496      518          L   0.93736   20       DESG.FLTIHVRWAMKNQYP     94.09
       497      519          A   0.96197   21      RKLAVTSMF.GPQHDENIWYC     96.40
       498      520          E   0.95973   20       LIKHTE.SDQVPAYRCGNFM     96.15
```
