## Supplemental Table 4 for "Oligomeric assemblies of plant biotin carboxylase revealed by cryo-EM and cross-linking"

Table S1. Summary of DSBU crosslinks in pennycress BC, in at least two sets of measurements.

|  |  | Band on SDS-PAGE | | | | Sample | | Fit into structure | |
| --- | --- | --- | --- | --- | --- | --- | --- | --- | --- |
|  |  | 100 kDa | | 200 kDa | | 1 mM DSBU replicate | |  |  |
|  | Pairs | pLink | Scout | pLink | Scout | R1 (pLink) | R2 (pLink) | Monomer 8HZ5.pdb | Dimer 8HZ5.pdb |
| 1 | 186 - 186 | X | X | X | X | X | X | N/A | No |
| 2 | 186 - 194 | X |  | X |  |  | X | Yes | - |
| 3 | 186 - 228 | X |  | X | X | X | X | Yes | - |
| 4 | 186 - 307 | X |  | X | X | X | X | Yes | - |
| 5 | 186 - 390 | X | X |  |  | X | X | No | No |
| 6 | 186 - 394 | X | X |  |  |  | X | No | No |
| 7 | 186– 416 | X | X |  |  |  |  | No | No |
| 8 | 194- 228 | X |  | X |  | X |  | Yes | - |
| 9 | 228 - 307 | X | X |  |  |  |  | Yes | - |
| 10 | 307 - 390 | X | X |  |  |  |  | Yes | - |
| 11 | 307– 390 | X |  |  |  | X | X | No | No |
| 12 | 307– 394 | X | X |  |  |  |  | No | No |
| 13 | 390– 390 | X | X |  |  | X | X | N/A | No |
| 14 | 390– 394 | X | X | X | X | X | X | Yes | - |
| 15 | 394- 416 | X | X |  |  |  |  | No | No |

Identified in both pLink and Scout

Identified in both replicates
