## Supplementary figures and images for "Oligomeric assemblies of plant biotin carboxylase revealed by cryo-EM and cross-linking"

### Video S2

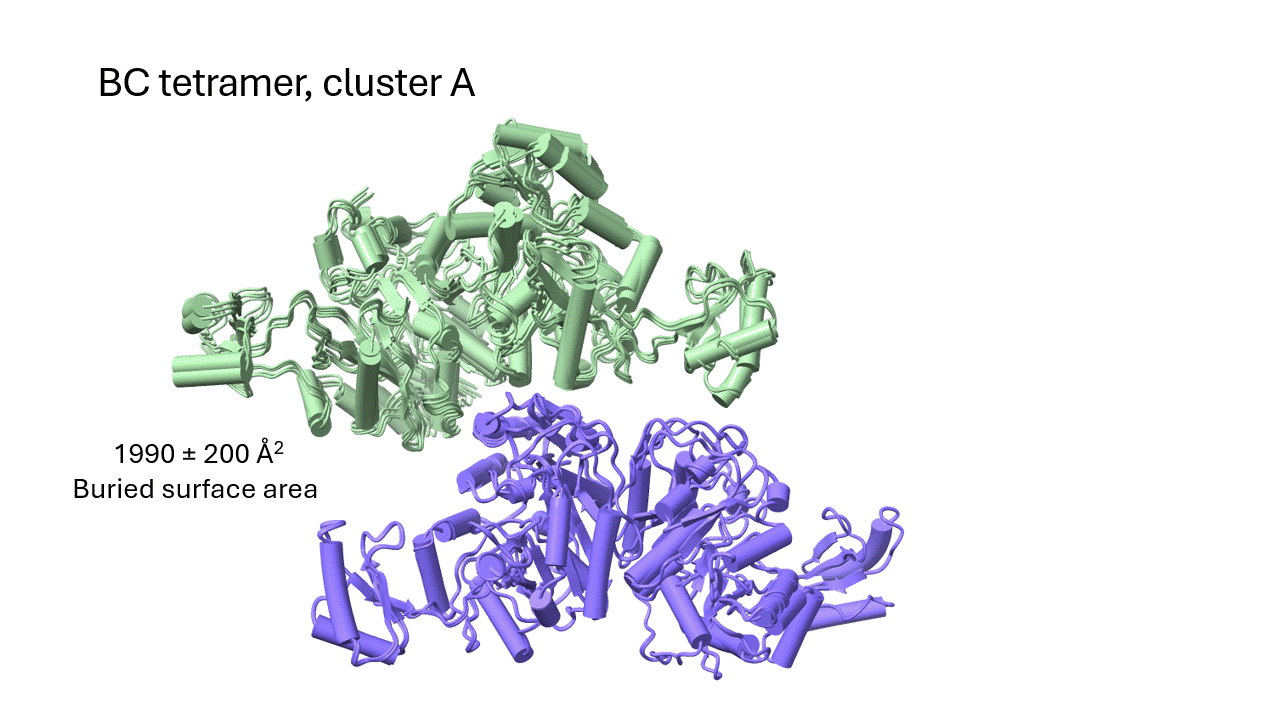
